# Organ-specific and conserved regulatory logic orchestrates gene expression in the embryonic mesothelium

**DOI:** 10.1101/2025.08.06.668886

**Authors:** Quang Minh Dang, Nicola Smart, Andia Nicole Redpath, Joaquim Miguel Vieira

**Affiliations:** Institute of Developmental and Regenerative Medicine, Department of Physiology, Anatomy & Genetics, University of Oxford, Oxford OX3 7TY, UK; High-Tech Center, Vinmec International Healthcare System, Hanoi, Vietnam; School of Cardiovascular & Metabolic Medicine & Sciences, King’s College London, London SE5 9NU, UK

## Abstract

The embryonic coelomic mesothelium undergoes epithelial-to-mesenchymal transition (EMT) to promote vascular growth and parenchymal development. A prominent example is the epicardium, which plays an essential role during heart development. Little is known about the mechanisms behind gene regulation in the coelomic mesothelium, or the organ-specific enhancer logic that endows specialization. Using gene regulatory network inference via multi-omic analysis, our study reveals *trans*- and *cis*-regulatory elements (CREs) that regulate mesothelial gene expression in three organs: heart, lung, and pancreas. We delineate pivotal transcription factors (TFs) and CREs specific to the epicardium, and show, in contrast, that the TF MAF orchestrates pan-mesothelial gene expression *via* conserved CREs, which are absent in non-mesothelial lineages. MAF may preserve mesothelial identity, evidenced by negative correlation with EMT and mouse-human conservation. Collectively, our work elucidates the regulatory logic behind cell type identity and leverages single-cell integration to gain insights into mammalian organ development.

**Highlights:**

- Coelomic mesothelia display distinct transcriptome linked to their morphogenic role
- Mesothelial markers associate with organ-specific *trans-* and *cis-*regulatory logic
- Pan-mesothelial gene regulatory mechanisms control canonical marker expression
- Integrating mouse and human datasets uncovers conserved dynamics of epicardial EMT

## Introduction

The mesothelium, an epithelium surrounding body cavities and visceral organs, serves as a critical interface between tissues. One of the best-known mesothelial tissues is the epicardium, which encases the heart. From embryonic day (E)11.5, epicardial cells undergo epithelial-to-mesenchymal transition (EMT) to form epicardium-derived cells (EPDCs). These mesenchymal derivatives invade the myocardium, differentiating into cardiac fibroblasts and mural cells. EPDCs thus provide cardiac precursors and, with the epicardium, supply paracrine signals that regulate myocardial growth, maturation, and coronary vessel development ^1^.

The coelomic mesothelium is also integral to the development of other organs it envelops, such as the lungs, liver, pancreas, and gonads, as reviewed elsewhere ^2^. Despite organ diversity, the mesothelium has shared features, including common genetic markers such as Wilms’ tumour 1 (*Wt1)*, which distinguish it from other lineages ^3–7^. Using these markers, lineage-tracing has revealed another common trait: multipotency. Mesothelial cells undergo EMT, generating diverse mesenchymal fates, including fibroblasts ^4, 8, 9^ and smooth muscle cells ^4, 10–12^. Consequently, *Wt1* knockout impairs EMT, causing significant defects in the heart and lungs ^3, 11, 13^. *Wt1* expression diminishes as development progresses and epicardial cells transition to a mesenchymal state ^14^, a pattern recapitulated in lung mesothelial cells ^4^.

Despite commonalities, mesothelial cells exhibit organ-specific differences. A key divergence is their origin. Most coelomic mesothelial cells arise from coelomic lining cells that acquire epithelial features ^2^, but the epicardium emerges from the proepicardial organ, a transient structure at the heart’s venous pole ^15^. Another divergence is the terminal fates of their mesenchymal derivatives. Mesothelial cells in chest cavity organs and the liver generate pericytes ^5, 10, 16–18^. However, pancreatic pericytes derive from a distinct *Nkx3.2*-expressing mesenchyme ^19^. Also, unlike in the liver, mesothelium-derived stellate cells in the pancreas are myofibroblast-like^7, 20^. Additionally, mesothelial cells also have organ-specific functions. In the pancreas, *Wt1*-positive mesothelial cells provide boundary separation from the stomach ^21^. Failure of mesothelial layer formation causes adhesion of the dorsal pancreas to the stomach, and systemic *Wt1* deletion leads to abnormal dorsal pancreas localization ^7^. Therefore, contrasting the pancreas with the heart and lungs, a prototypical mesothelium-encased organ, can reveal the full spectrum of mesothelial behaviour.

These observations underscore the coelomic mesothelium’s importance as a source of organ-specific cell types and regulatory signals that shape visceral morphogenesis. Deciphering the underlying gene regulatory networks (GRNs) is key to understanding mesothelial identity and diversification. GRNs, which map interactions between transcription factors (TFs) and cis-regulatory elements (CREs) including enhancers, control gene expression and cellular identity ^22, 23^. A GRN-centric analysis is therefore central to understanding the mesothelium’s cellular plasticity.

Here, we performed ATAC-seq of the *in vivo* epicardium at E11.5 and E17.5 and *in vivo* EPDCs at E13.5. We integrated these data with available RNA-seq, ATAC-seq, and CUT&RUN-seq datasets to profile the GRNs governing mesothelial spatiotemporal behaviour in the heart, lung, and pancreas. We identify that specific CREs and TFs likely control spatiotemporal mesothelial expression and highlight their potential relevance to epicardial EMT. We also documented conserved, pan-mesothelial regulatory elements and mechanisms absent from non-mesothelial lineages. We identified several such elements driving the spatiotemporal expression of cytokeratins, linked to MAF. By curating an extensive scRNA-seq dataset of murine and human epicardial cells and an integrated scRNA-seq dataset profiling murine epicardial EMT, we reinforced MAF’s human relevance and its importance for mesothelial identity, noting its repression as epicardial cells undergo EMT. By pinpointing pan-mesothelial regulatory mechanisms, our research opens new avenues for therapeutic strategies leveraging the coelomic mesothelium’s regenerative capacity.

## Results

### Single-cell genomics for benchmarking mesothelial cells across embryonic organs and reappraisal of the role of cadherins in EMT

To understand the GRNs governing mesothelial cell identity, we profiled gene expression and chromatin accessibility in the mouse heart, lung, and pancreas at E13.5, a stage of active EMT in all three organ linings (Figure 1A) ^4, 7, 24^. For the epicardium, we used our scRNA-seq ^14^ and bulk ATAC-seq data from FACS sorted cells ^25^ (Figures 1B, S1A). For lung and pancreas, we examined public scRNA- and scATAC-seq datasets (Figures 1C, 1D, S1B, S1C) ^26^. We confirmed *Wt1* and Uroplakin 3B (*Upk3b*) as robust markers for mesothelial clusters in all three organs and data modalities (Figure 1E-G, S1A–S1C).

**Figure 1.**
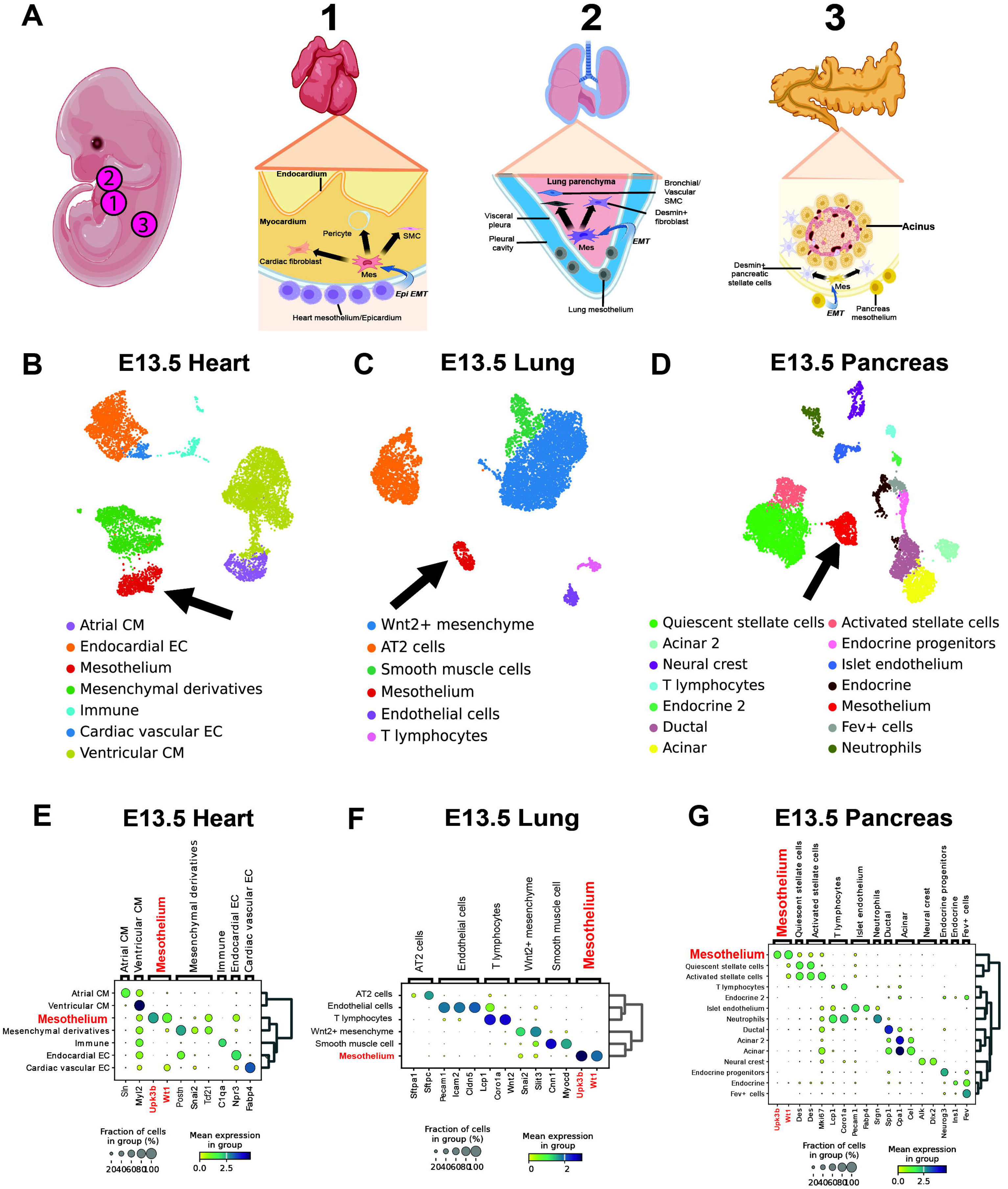
Multi-omic profiling of the embryonic mesothelia across organs. (A) Schematic illustrating cellular contributions of the coelomic mesothelium to the embryonic heart (1), lung (2), and pancreas (3) via epithelial-to-mesenchymal transition (EMT). Created with Biorender.com (B) Uniform manifold approximation and projection (UMAP) of the E13.5 mouse heart (7,867 cells) based on gene expression. Black arrow indicates the epicardium cluster. (C) UMAP of the E13.5 mouse lung (5,925 cells). (D) UMAP of the E13.5 mouse pancreas (7,113 cells). (E) Dot-plot of marker gene expression in the E13.5 heart. The mesothelium was identified by *Upk3b* and *Wt1* expression. Dot size represents the fraction of cells expressing each gene. (F) Dot-plot of marker gene expression in the E13.5 lung. (G) Dot-plot of marker gene expression in the E13.5 pancreas. *AT2: Alveolar-type 2; CM: Cardiomyocyte; EC: Endothelial cell.* **See also Figure S1**

A major roadblock to understanding mesothelial cell identity and function is the paucity of knowledge of cadherin activity. Cadherins are critical for EMT by forming adherens junctions and mediating cell-cell adhesion, therefore maintaining epithelial identity ^27, 28^, but their specific roles in the coelomic mesothelium are contested and unclear. The previously implicated role of CDH1 (E-cadherin) in epicardial EMT has been refuted ^3, 13^. We examined cadherin expression and found that only N-cadherin (*Cdh2*), P-cadherin (*Cdh3*), and cadherin-11 (*Cdh11*) were notably expressed across all mouse organ mesothelia (Figure S1D). Notably, E-cadherin (*Cdh1*) is absent from all organ mesothelia at E13.5, consistent with other epicardial studies ^3^, suggesting it is not a primary target of EMT-inducing transcription factors in the mesothelium.

To further delineate cadherins’ roles in EMT, we devised a computational approach leveraging single-cell genomics and lineage tracing to reconstruct the mesothelial EMT trajectory *in silico*, focusing on the heart due to data availability (see Methods, Figure S1E). We integrated our published data with other datasets, creating a 7,332-cell dataset spanning E12.5 to E17.5 (Figure S1F). This integrated dataset includes epicardial cells, their mesenchymal progeny, and differentiated cells (Figure S1F, S1G). Pseudotime scoring predicted branched pathways with distinct fates (epicardial maturation, or differentiation into smooth muscle cell progenitor, cardiac fibroblasts, pericytes, or valvular interstitial cells). The integrated dataset confirms *Cdh1* is absent across the epicardial lineage (Figure S1H). *Cdh2* expression positively correlates with the EMT trajectory (Figure S1H, S1I), supporting its requirement for epicardial migration into the myocardium – a foundational step for EMT ^29^. Conversely, the highly expressed cadherins *Cdh3* and *Cdh11* are downregulated along the trajectories (Figure S1H, S1J, S1K), suggesting they may act as guardians of epicardial identity.

### Organ mesothelia exhibit transcriptional and epigenomic concordance

To reveal the epicardial GRN, we analyzed gene expression and chromatin accessibility (Figure S1E). We used our E13.5 ATAC-seq data ^25^ and identified high-quality peaks through an iterative filtering approach ^30^. We validated our bulk ATAC-seq dataset (dataset 1) against published E13.5 scATAC-seq data (dataset 2) ^26^, the only other dataset capturing epicardial cells at this stage, to our knowledge. We subsetted the *Upk3b*-expressing epicardial cluster from dataset 2 (Figure 2A) and found a strong correlation between datasets (Figure 2B, 2C), with dataset 1 capturing 86.6% of peaks in dataset 2. The 81,067 peaks unique to dataset 1 likely reflect the higher number of epicardial cells sampled (∼10,000-50,000 vs. 649), capturing more chromatin features. We further leveraged H3K4me3 and H3K27ac CUT&RUN-seq data from cultured murine embryonic epicardial cells (MEC1) to identify active promoters and enhancers ^25, 31^ (Figure S2A, S2B). Many of these regions were accessible in dataset 2 (18058 enhancers and 16514 promoters; Figure 2D) and other organ mesothelia (23704 enhancers and 14550 promoters; Figure S2C, S2D). To further demonstrate data quality, a weighted gene co-expression analysis identified organ-specific mesothelial gene modules (Figure S2E) and GO term analysis identified relevant terms such as “heart morphogenesis” and “lung development” (Figure S2F). All transcriptional modules were of high quality and conserved in the ATAC-seq datasets as determined by the Z-statistics metric ^32^ (Figure S2G).

**Figure 2.**
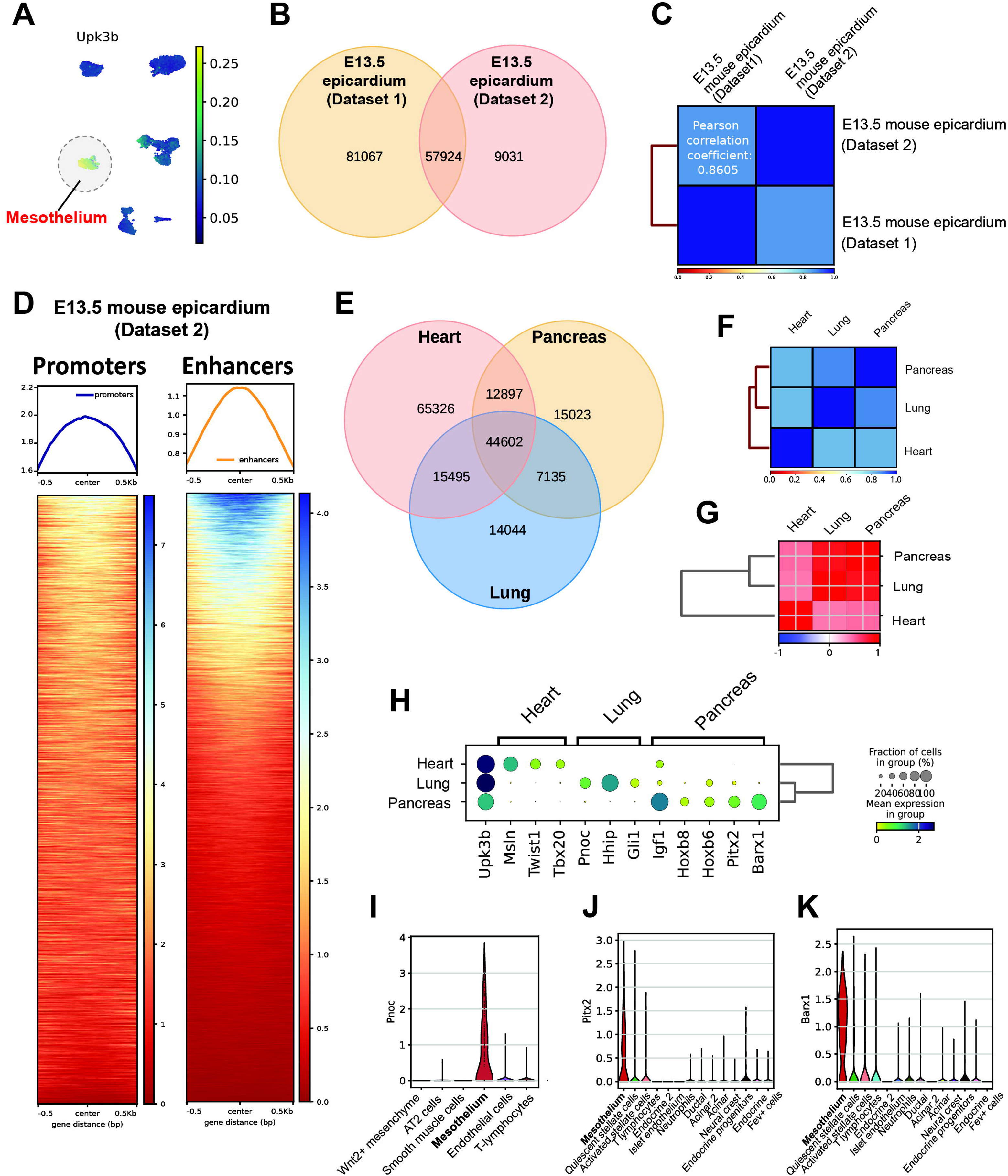
Transcriptomic and epigenomic concordance of the coelomic mesothelium across organs. (A) UMAP of the E13.5 mouse heart scATAC-seq dataset (7,292 cells). Epicardial cells (649 cells) are highlighted, identified by enriched *Upk3b* gene activity. (B) Venn diagram of peak intersection from the E13.5 epicardium across two studies. Dataset 1 is from ^25^; dataset 2 is from ^26^. (C) Correlation matrix for epicardial ATAC-seq from two studies based on accessible peak signatures. Pearson’s correlation coefficient shown and.strong correlation is indicated in blue. (D) Heatmap of normalized peak scores of epicardial ATAC-seq profiles ^98^ in active epicardial promoter and enhancer regions. Active enhancers and promoters were identified from H3K27ac and H3K4me3 CUT&RUN-seq of the MEC1 epicardial cell line ^25^. (E) Venn diagram of ATAC-seq peak intersection from the E13.5 mesothelium across the heart, lung, and pancreas. (F) Correlation matrix for E13.5 organ mesothelia based on chromatin accessibility signatures. (G) Correlation matrix for E13.5 organ mesothelia based on transcriptomic signatures. (H) Matrix dot-plot showing mean expression of enriched marker genes for each E13.5 mesothelium. (I) Violin plot of normalised *Pnoc* expression in the E13.5 mouse lung. (J) Violin plot of normalised *Pitx2* expression in the E13.5 mouse pancreas. (K) Violin plot of normalised *Barx1* expression in the E13.5 mouse pancreas. **See also Figure S2**

We then compared other organ mesothelia. ATAC-seq peak intersection revealed conserved chromatin accessibility, with 54.9% and 56.0% of peaks in lung and pancreas mesothelium, respectively, being accessible in all three (Figure 2E), which we term consensus mesothelial elements. Epigenomic and transcriptomic concordance between organ mesothelia is strong (Figure 2F, 2G). However, the correlation with the epicardium is weaker, with 47.2% of its ATAC-seq peaks being unique, alluding to organ specificity (Figure 2E). A metacell-based differential gene expression analysis identified enriched markers: Mesothelin (*Msln*), *Tbx20*, and *Twist1* for the epicardium; Hedgehog interacting protein (*Hhip*) and *Gli1* for the lung mesothelium; and Insulin growth factor 1 (*Igf1*) and Hox TF family members *Hoxb6* and *Hoxb8* for the pancreas mesothelium (Figure 2H, Supplementary table 1,2). *Msln*’s epicardium-specificity at E13.5 is intriguing, as it’s expressed later in other mesothelia ^33, 34^. Among these markers, we identified *Pnoc* (Figure 2I), *Pitx2* (Figure 2J), and *Barx1* (Figure 2K) as distinguishing mesothelial cells within their respective organs. The mesothelium-restricted expression of *Pitx2* and *Barx1* is consistent with previous findings ^21, 33^, and their chromatin-inferred gene activities reliably distinguish the pancreas mesothelium (Figure S2H, S2I). Conversely, *Pnoc* has not been previously described as a lung mesothelium marker.

### Inference of mesothelial GRNs highlights transcriptional regulons as putative encoders of mesothelial identity across organs

To understand the regulatory code underpinning mesothelial identity and function, we inferred GRNs using the SCENIC+ protocol, which predicts TF-enhancer-gene (triplet) relationships as regulons ^23^. We created a custom motif database by combining databases from SCENIC+ and Yi et al. ^35^, with the latter included to enhance mouse regulon discovery. For high-confidence predictions, we selected top importance scores for each triplet (Supplementary table 3,4). We cross-validated putative epicardial enhancers using MEC1 H3K27ac CUT&RUN-seq data and corroborated predicted E-P connections with the Activity-By-Contact (ABC) model ^36^. We only considered regulatory relationships predicted by both SCENIC+ and the ABC model in the epicardium. For non-cardiac mesothelia lacking H3K27ac data, which can degrade the ABC model’s performance, we relied only on SCENIC+.

Contrasting organ mesothelial GRNs, we identified regulons with enriched activity in the epicardium, including TBX20, which exhibits strong epicardial specificity (Figure 3A). In the lung mesothelium, regulons for Fox family members, particularly FOXF1 and FOXP2, were specific (Figure 3B), consistent with their roles in lung morphogenesis ^37, 38^. Genome-wide footprinting confirmed differential DNA binding for FOXF1 and FOXP2 between the epicardium and lung mesothelium (Figure 3D). For the pancreas mesothelium, ISL1 and *PITX2* regulons were central, showing cell-type specificity (Figure 3C) and enhanced TF binding compared to the epicardium (Figure 3E). ISL1 is known for its prominent role in pancreas morphogenesis ^39, 40^, being required for endocrine islet cells differentiation.

**Figure 3.**
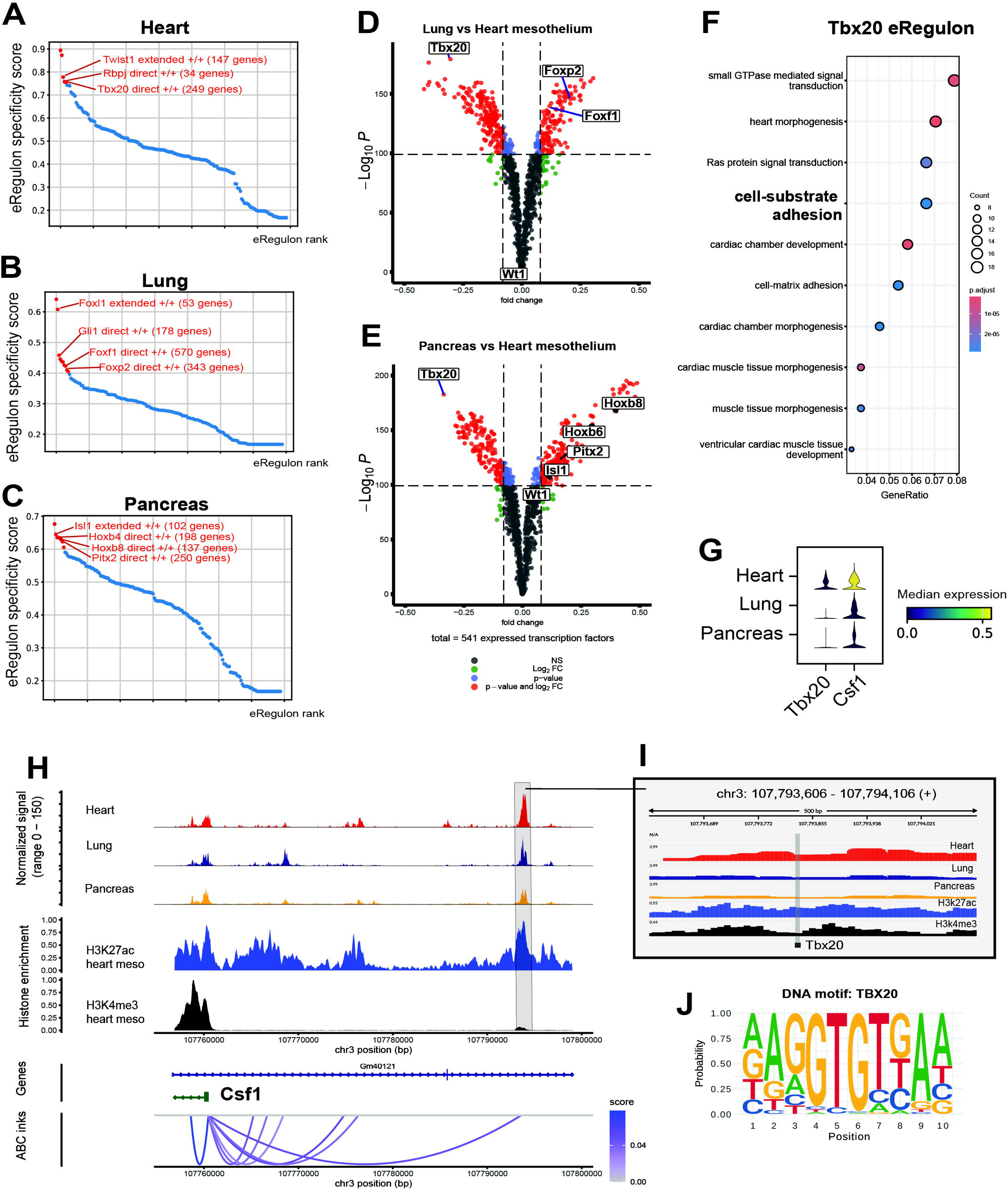
Mesothelial gene expression is powered by organ-specific gene regulatory networks. (A) Top activator regulons in the heart mesothelium ranked by cell-type specificity. Top 10 specific regulons are highlighted in red. (B) Top activator regulons in the lung mesothelium ranked by cell-type specificity. (C) Top activator regulons in the pancreas mesothelium ranked by cell-type specificity. (D) Volcano plot of differential TF binding activity comparing the heart and lung mesothelium. Differential binding activity against the −log10(p value) of all investigated TF motifs is shown; each dot represents one motif. Only TFs expressed in at least one organ mesothelium are retained. Significantly differential motifs are coloured in red (FC>0.08, - log_10_P > 100) (E) Volcano plot of differential TF binding activity comparing the heart and pancreas mesothelium. (F) GO term over-representation of TBX20 regulon target genes. (G) Stacked violin plot showing median expression of *Tbx20* and *Csf1* in E13.5 organ mesothelial cells. (H) Genome tracks showing a distal CRE, a putative enhancer (highlighted), regulated by TBX20, and predicted to regulate the *Csf1* promoter. Merged normalised ATAC-seq tracks displayed. Merged histone tracks show MEC1 epicardial H3K27ac (N = 3) and H3K4me3 (N = 4) CPM-normalised scores. The link plot shows predicted enhancer-promoter (E-P) connections in the epicardium; link colour denotes the ABC model score (the confidence of E-P prediction). (I) Footprints of the *Csf1* enhancer highlighted in (H) across organ mesothelia. The position of TOBIAS predicted TBX20 binding site is highlighted. (J) The TBX20 mouse motif logo. **See also Figure S3, S4, S5**

We studied the predicted GRNs in greater detail, focusing on highly specific regulons with many target genes. In the epicardium, TBX20 has the largest number of predicted targets. While known to be indispensable for cardiomyocyte biology ^41–43^, its role in the epicardium is unstudied. We found epicardium-enriched DNA binding activity for TBX20 (Figure 3D, 3E). GO analysis of its target genes revealed enriched terms relevant to cardiac morphogenesis and, notably, “cell-substrate adhesion”, suggesting unique epicardial functions (Figure 3F, Supplementary Table 5). A gene comprising this term is Macrophage Colony-Stimulating Factor (*Csf1*), a cytokine that regulates cardiac macrophages ^44^. *Csf1* was enriched in the epicardium (Figure 3G), consistent with its role in yolk-sac derived macrophage recruitment and cardiac regeneration ^45–48^. We identified a CRE (chr3:107793606-107794106) with a TBX20 binding site that has enriched accessibility in the epicardium (Figure 3H). Footprinting showed the TBX20 motif is bound only in the epicardium (Figure 3I, 3J). The region is an active enhancer (H3K27ac mark) and was identified by the ABC model as a *Csf1* regulator (Figure 3H). We thus conclude that TBX20 may be a regulator of *Csf1*.

Using similar strategies, we found other examples of TBX20-driven, epicardium-specific regulation of *Fgf2*, *Sema3d,* and *Rarres2*, genes critical for epicardial function (Figure S3A-I, Supplemental table 3) ^15, 25, 49^. TBX20 may also contribute to epicardial EMT through genes regulating cell adhesion. We identified TBX20-mediated regulation of plakophilin-2 (*Pkp2*), a desmosome component expressed mostly only in the epicardium (Figure S3J), potentially reflecting the unique mechanical stress experienced by cardiac tissue. We found two putative epicardium-accessible enhancers (chr16:16219711-16220211 and chr16:256606-16257106) bound by TBX20 linked to *Pkp2* (Figure S3K). Given the role of desmosomes in maintaining cellular adhesion of the mesothelial layer, their disruption may correlate with epicardial EMT. We reason that *Pkp2* expression will change over the course of EMT, and so will the activity of its enhancers. Therefore, we tested the activity of our proposed CREs as an additional strategy to validate their credibility. We generated ATAC-seq data that profiled the *in vivo* EPDCs at E13.5, when EMT peaks. We utilised the inducible *Wt1*^CreERT2^ mouse, crossed with R26R-tdTomato (tdTom) reporter, to trace the epicardial lineage as previously described ^14^. Due to their negative surface expression of podoplanin (PDPN) ^25, 50^, EPDCs were sorted as CD31-tdTom+PDPN-cells for bulk ATAC-seq, and systematically compared against E13.5 epicardial ATAC-seq data ^25^. Both enhancers had significantly lower chromatin accessibility in EPDCs (enhancer in S3L: Log2FC −1.15 and FDR 2.23*10-5; enhancer in S3M: Log2FC −2.10 and FDR 1.34*10-6) (Supplementary table 6), aligning with the downregulation of *Pkp2* during EMT (Figure S3N).

TBX20 may also regulate epicardial layer integrity via adherens junctions, which in turn involves cadherins. *Cdh2* (N-cadherin) expression increases during EMT (Figure S1I). We propose this is enabled by TBX20 regulation through a distal enhancer (Figure S3O, Supplemental table 3) that exhibits slightly increased (17% higher) accessibility in EPDCs (Supplementary table 6). Collectively, TBX20 may direct transcriptional programs that provide cardiac-specific context to the epicardial GRN and are linked to EMT.

### Master transcriptional regulators may control lung and pancreas mesothelial marker genes via distal enhancers

We then studied prominent regulons in the lung mesothelium. One of the top lung-specific regulons *GLI1*, the Hedgehog signaling effector indispensable for mesothelial cell invasion ^4^ showed enriched expression in lung mesenchyme (Figure S4A). Our analysis suggests *Hhip* is a target gene of GLI1 (Figure S4B), confirming its known dependence on GLI1 ^51^.

Another key lung-specific regulon is FOXF1 (Figure 3B), which is essential for lung morphogenesis ^37^ and not expressed in non-lung mesothelia (Figure S4C). Our findings indicate FOXF1 may regulate *Pnoc*, a lung mesothelium marker (Figure 2I), via a lung-specific CRE (Figure S4D). Another putative target is FOXP2, essential for lung epithelial development ^38^. We found *Foxp2* is also highly expressed in the lung mesothelium (Figure S4E), possibly modulated by FOXF1 binding to its promoter (Figure S4F). FOXF1 may also regulate *Gli1* itself via a distal CRE and TBX2, which contributes majorly to lung development and mesothelial lineage ^52–54^, via a lung-specific CRE (Figure S4G, S4H).

In the pancreas mesothelium, we identified key regulons that may drive marker gene expression (Figure 2H). A prominent example is BARX1, which may regulate *Hoxb8* via a promoter-proximal CRE (Figure S5A). BARX1-deficient mice have spleen defects and abnormal pancreas location, suggesting a role for BARX1 in pancreatic morphogenesis ^55^. Another key regulon is *PITX2*, known for left-right patterning function ^56^ but predominantly expressed in the E13.5 mesothelium (Figure 2J). We found a distal region predicted to regulate *Igf1* via PITX2 (Figure S5B). IGF1 plays a significant role in exocrine pancreas development by acting through its receptor IGF1R ^57^. Another PITX2 target is the Cadherin 6 (*Cdh6*) promoter (Figure S5C), uniquely expressed in the pancreas mesothelium (Figure S1D). CDH6, although unstudied in the pancreas, may be critical for maintaining epithelial identity, judging by its involvement in forming the kidney epithelium ^58^ through formation of adherens junctions ^59^. Lastly, the ISL1 regulon is predicted to regulate *Gcg*, a gluconeogenesis hormone, via a CRE with three ISL1 binding sites (Figure S5D, S5E).

### Spatiotemporal gene expression of the coelomic mesothelium is conserved and is linked to the activities of *cis*-regulatory elements

We have so far described organ specifications at a single time point. However, expression can be temporal, as highlighted by *Msln*, which is expressed in the epicardium at E13.5 but appears later in other mesothelia ^33, 60^. *Msln* is an important marker distinguishing mesothelial cells from other lineages across organs ^34, 61^, over lifetime ^34^, and holds significant roles in mesothelioma biology ^62–64^. Using *Msln* as a case study, we elucidated principles of temporal gene regulation in the mesothelium.

We scanned the *Msln* promoter for TF motifs and found candidates including RBPJ, SOX proteins, TCF proteins, and AP-1 family members such as NRF2/NFE2l2 (Figure S6A). The promoter is accessible only in the epicardium (Figure 4A, 4B). However, two proximal CREs, marked by H3K27ac, are also linked to the *Msln* promoter (Figure 4B). Both CREs contain WT1 binding sites (Figures 4C, 4D, S6B), suggesting WT1 is a direct regulator. Since *Msln* is majorly downregulated during EMT (Figure S6C), we used our EPDC ATAC-seq to show these CREs are also inactive in mesenchymal derivatives (Figure S6D).

**Figure 4.**
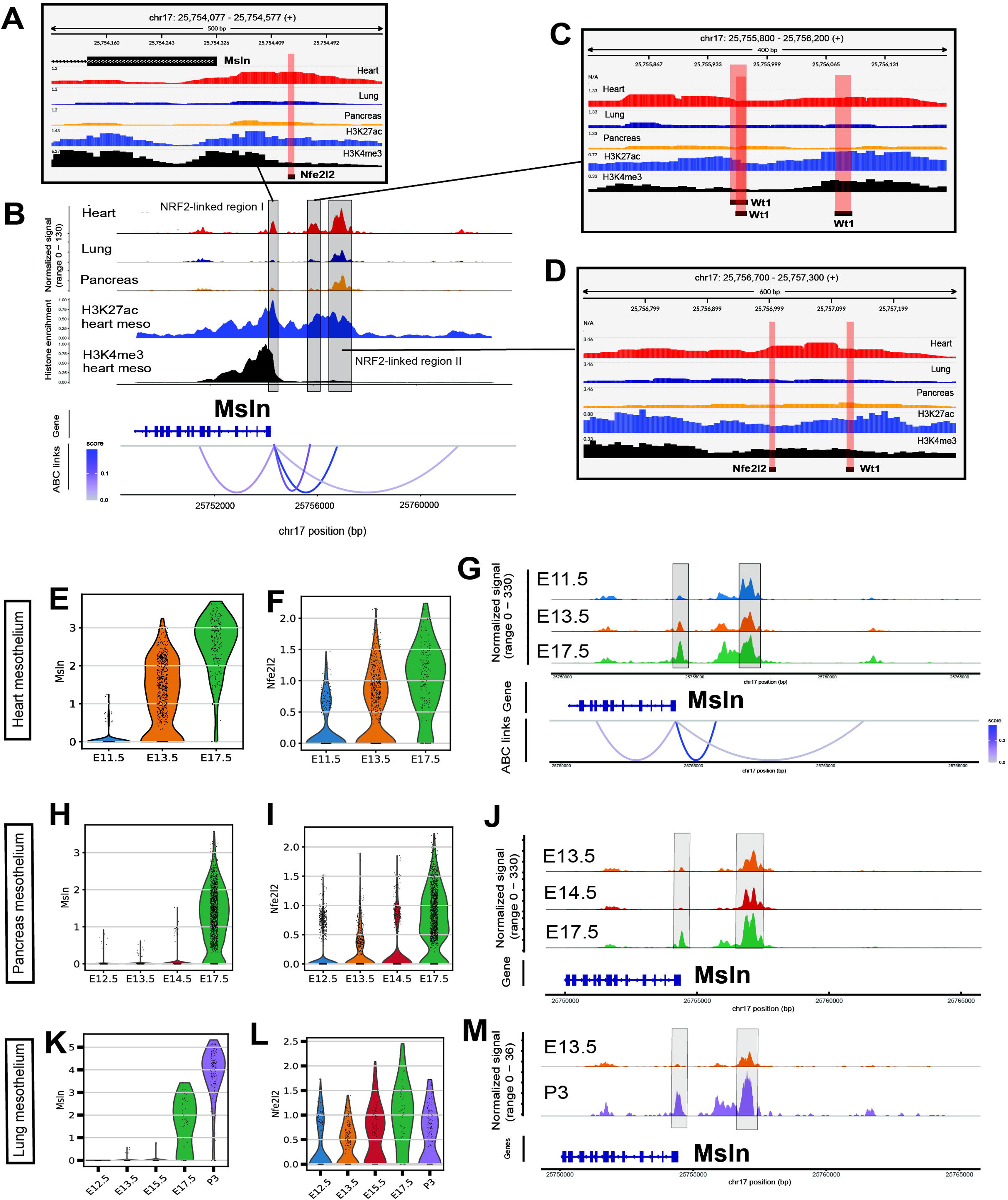
Spatiotemporal regulation of *Msln* is conserved across organ coelomic mesothelia. (A) Footprints of the *Msln* promoter. The NRF2/NFE2L2 motif is highlighted. (B) Genome tracks of the *Msln* locus. The promoter and two distal CREs are highlighted. (C) Footprints of the CRE (chr17:25755800-25756200) linked to the *Msln* promoter. WT1 motifs are shown. (D) Footprints of the CRE (chr17:25756700-25757300). WT1 and NRF2/NFE2L2 motifs are shown. (E) Violin plot of normalised *Msln* expression in the mouse epicardium at E11.5, E13.5, and E17.5. (F) Violin plot of normalised *Nfe2l2* expression in the mouse epicardium at E11.5, E13.5, and E17.5. (G) Genome tracks of the *Msln* locus in epicardial cells at E11.5, E13.5, and E17.5. The promoter is highlighted. Merged normalised ATAC-seq tracks displayed (N = 3). The link plot shows ABC model predicted enhancer-promoter (E-P) connections from all three stages. (H) Violin plot of normalised *Msln* expression in the mouse pancreas mesothelium at E12.5, E13.5, E14.5 and E17.5. (I) Violin plot of normalised *Nfe2l2* expression in the mouse pancreas mesothelium at E12.5, E13.5, E14.5 andE17.5. (J) Genome tracks of the *Msln* locus in pancreas mesothelial cells at E13.5, E14.5, and E17.5. NRF2-linked regions are highlighted. All tracks were pseudo-bulked and TF-IDF-normalised. (K) Violin plot of normalised *Msln* expression in the mouse lung mesothelium at E12.5, E13.5, E17.5, and P3. (L) Violin plot of normalised *Nfe2l2* expression in the mouse lung mesothelium at E12.5, E13.5, E17.5, and P3. (M) Genome tracks of the *Msln* locus in lung mesothelial cells at E13.5 and P3. NRF2-linked regions are highlighted. **See also Figure S6**

To test if these CREs regulate *Msln* expression over time, we generated bulk ATAC-seq datasets of the *in vivo* epicardium at E11.5 and E17.5. *Msln* expression increased as the epicardium matures (Figure 4E), consistent with protein levels ^65^. Mirroring this, chromatin accessibility of CREs increased with development, peaking at E17.5 (Figure 4G, Supplementary table 9, 10). SCENIC+ analysis of our temporal data identified NRF2 as the highest confidence candidate TF (Supplementary table 11). NRF2 has binding sites in the *Msln* promoter and one enhancer (Figure 4A, 4D), and its expression (*Nfe2l2*) increases throughout development, correlating with *Msln* (Figure 4F). We now label these regions as NRF2-linked regions (Figure 4B, Figure S6D).

We tested if these NRF2-linked regions also regulate *Msln*’s temporal expression in other organs. In the pancreas, *Msln* and *Nfe2l2* expression becomes prominent from E17.5 (Figure 4H, 4I). Consistently, ATAC-seq shows NRF2-linked regions have elevated accessibility at E17.5 (Figure 4J). This pattern of accessibility and expression is conserved in the lung mesothelium (Figure 4K, 4L). Here, for the lung, we used postnatal day 3 (P3) ATAC-seq data due to a lack of high-quality data capturing the mesothelium in late embryonic development (between E13.5 and P3). This suggests a conserved mechanism of temporal gene expression, albeit with variation in developmental timing.

Our temporal data also reinforced other CRE predictions we have made. In the epicardium, the expression of TBX20 and its targets *Fgf2* and *Csf1* decreases from E13.5 to E17.5 (Figure S6E), as does the accessibility of their nominated CREs (comparing E17.5 against E13.5: *Csf1* CRE: Log2FC −0.64, FDR 5.39*10^−14^; *Fgf2* CRE: Log2FC −0.81, FDR 1.15*10^−10^; Figure S6F, S6G). In the pancreas, expression of BARX1 and its target gene correlates over time (Figure S6H), mirrored by the accessibility of the *Hoxb8*-linked CRE (Figure S6I). A similar correlation was seen for ISL1/*Gcg* (Figure S6J, S6K). Collectively, these examples illustrate GRN-dependent temporal gene regulation in the mesothelium.

### Comparison against other cardiac cell types reveals mesothelial-specific *cis*-regulatory grammar and regulons

We next explored the regulatory dynamics that distinguish the coelomic mesothelium from other cell lineages, which may reveal master regulators of mesothelial identity. Using the heart as an example, we selected ventricular cardiomyocytes (CM) and cardiac vascular endothelial cells (EC) for comparison (Figure 5A). Using markers *Myl2* and *Fabp4*, we obtained transcriptomic and chromatin profiling of E13.5 CM and vascular EC (Figures 5B, S7A, S7B). As expected, epicardial cells are more similar to other organ mesothelia than to other cardiac cell types, based on chromatin accessibility and transcriptomics (Figure S7C-E). Combined analysis with cardiac lineages reveals enriched expression of previous markers that distinguish organ mesothelia (Figure S7F).

**Figure 5.**
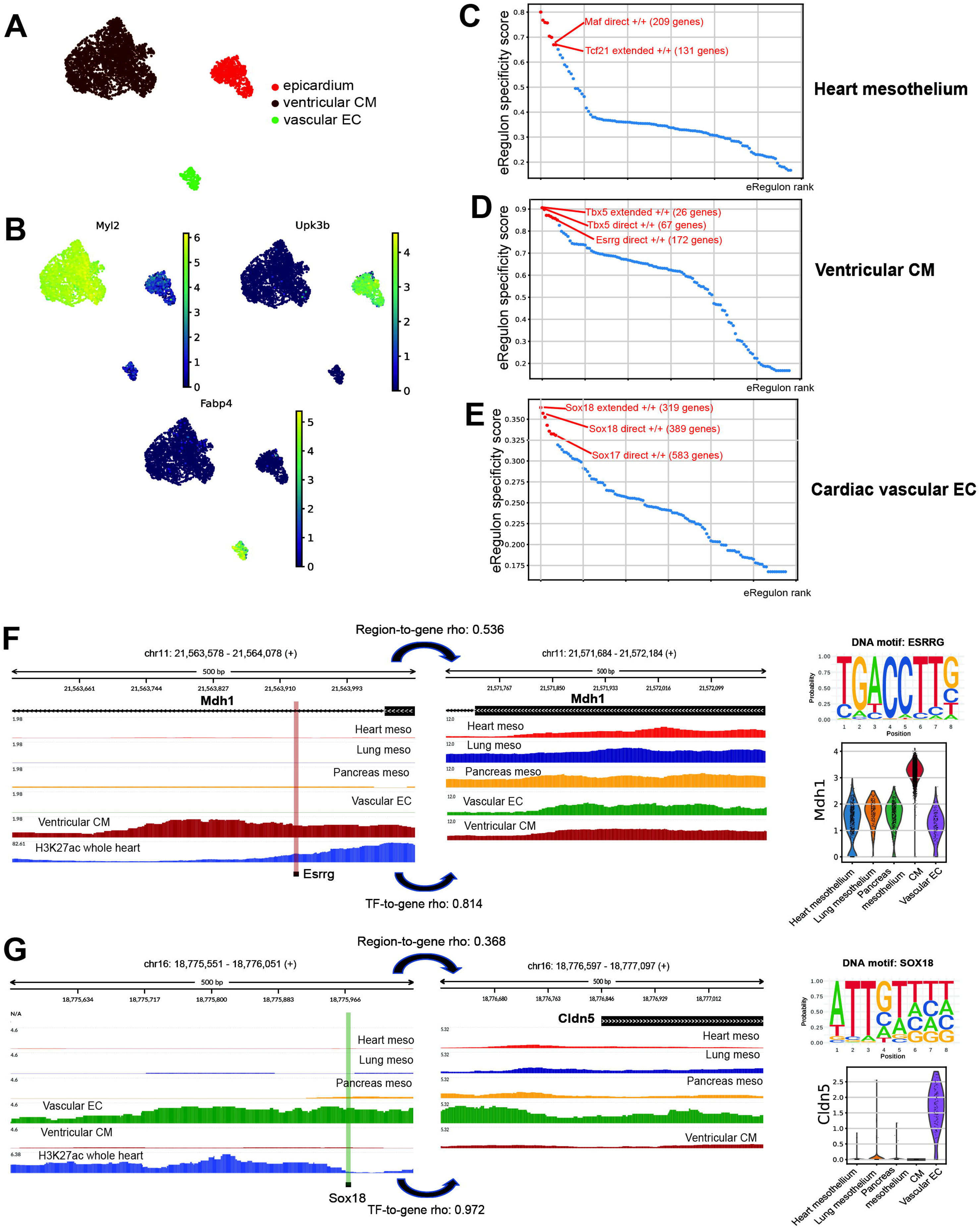
Comparison with other cardiac cell types reveal mesothelial-specific gene regulation landscape. (A) UMAP of E13.5 mouse cardiac cell types (3,700 cells). (B) UMAP feature plots of cardiac cell type markers: *Myl2* (CM), *Upk3b* (heart mesothelium), Fabp4 (vascular EC).(C) Top activator regulons in the epicardium ranked by cell-type specificity. Top 10 specific regulons are highlighted in red.(D) Top activator regulons in CM ranked by cell-type specificity. (E) Top activator regulons in vascular EC ranked by cell-type specificity. (F) Genome tracks showing an ESRRG-regulated CRE predicted to regulate *Mdh1*. The violin plot shows normalised *Mdh1* expression. (G) Genome tracks showing a SOX18-regulated CRE predicted to regulate *Cldn5*. The violin plot shows normalised *Cldn5* expression. *CM: Ventricular cardiomyocytes; EC: Endothelial cells* **See also Figure S7**

GRN inference highlighted cell-type-specific regulons, including MAF and TCF21 for the epicardium (Figure 5C), ESRRG and TBX5 for CMs (Figure 5D), and SOX17 and SOX18 for vascular ECs (Figure 5E). These TFs are known regulators of their respective cell types, with ESRRG and TBX5 being required for CM maturation, while SOX17 and SOX18 are regulators of the endothelial/hematopoietic lineage commitment ^66–69^. Differential footprinting analysis using curated TF motifs (Figure S7G) confirmed their cell-type-specific binding and the epicardium-enriched binding of MAF and TCF21 (Figure S7H, S7I). Examining these regulons, we identified validated TF-gene relationships, such as ESRRG regulating *Mdh1* in CMs ^70^ (Figure 5F) and SOX18 regulating *Cldn5* in vascular ECs ^71^ (Figure 5G). The strength of our inference, demonstrated in other cell types, reinforces our subsequent findings on candidate TFs that distinguish mesothelial cells from other cardiac lineages.

### Evolutionarily conserved *cis*-regulatory elements drive mesothelial gene expression patterns shared across organs

Based on the cardiac lineage comparison, we studied the regulatory targets of the epicardial-specific regulons MAF and TCF21. TCF21 is highly expressed in the epicardium and required for EPDC commitment to a fibroblast fate ^72, 73^. MAF, on the other hand, has not been studied in coelomic mesothelial cells. Both TFs have limited expression in CMs and CECs (Figure S8A). MAF, in particular, is expressed in both cardiac and non-cardiac mesothelia.

MAF’s regulon revealed its control of cytokeratins (Krt), which form intermediate filaments that facilitate epithelial cell adhesion by interacting with desmosomes ^74^. Several cytokeratins, including *Krt7*, *Krt8*, and *Krt18,* are epicardial markers ^61, 75^. We provide evidence that MAF regulates *Krt8* and *Krt18* via putative enhancers conserved across multiple organ mesothelia. *Krt18* showed pan-mesothelial expression (Figure 6A), and we identified a shared accessible enhancer (chr5:102016032-102016532) linked to its promoter (Figure 6B). As cytokeratins are critical for intercellular adhesion and thus may be inactive after EMT, we confirmed this CRE has significantly lower accessibility (Log2FC: −2.04, FDR: 4.67*10-6) in EPDCs (Figure 6C). A similar logic was applied to *Krt8*, another pan-mesothelial cytokeratin (Figure 6D), which has a conserved CRE (chr15:102005234-102005734) (Figure 6E) with reduced accessibility in EPDCs (Log2FC: −1.79, FDR: 5.88*10-7; Figure 6F). The reduced CRE activity correlates with the downregulation of *Krt8*, *Krt18*, and *Maf* during epicardial EMT (Figure 6G), reinforcing MAF as a regulator of cytokeratins and mesothelial identity.

**Figure 6.**
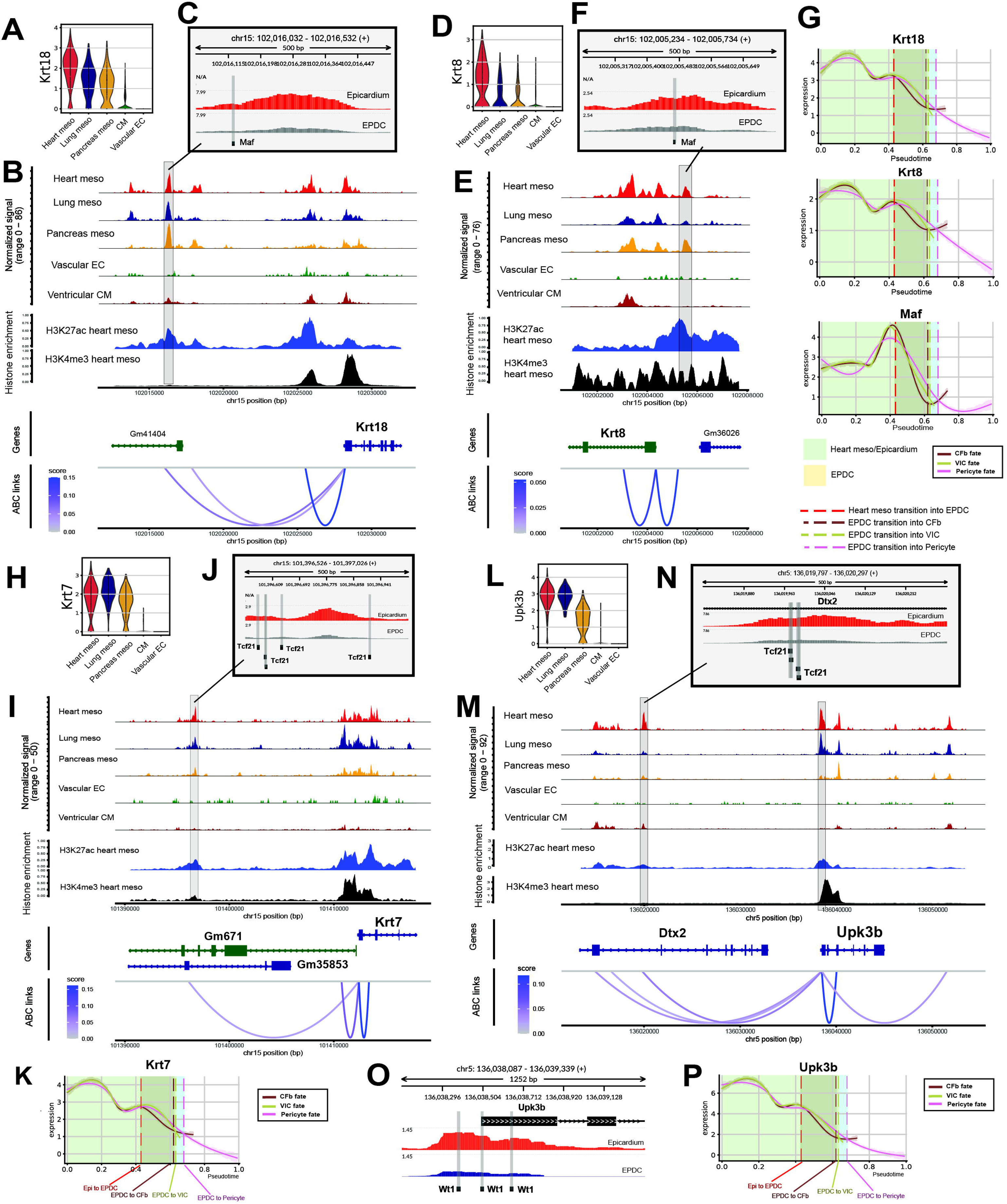
Conserved CREs regulate mesothelial gene expression. (A) Violin plot of normalised *Krt18* expression in mesothelia, CM, and Vascular EC. (B) Genome track of the *Krt18* locus showing a putative enhancer (highlighted). (C) Normalised ATAC-seq signals of the CRE from (B) in FACS sorted epicardial cells and EPDCs. ATAC-seq read counts were normalised using the trimmed mean of M values (TMM) method. The MAF motif is shown. (D) Violin plot of normalised *Krt8* expression in mesothelia, CM, and CECs. (E) Genome track of the *Krt8* locus showing a putative enhancer (highlighted). (F) Normalised ATAC-seq signals of the CRE from (E) in FACS sorted epicardial cells and EPDCs. Position of MAF binding site in the enhancer is shown. (G) Imputed expression of *Maf*, *Krt8*, and *Krt18* along the epicardial EMT pseudotime trajectory. Three trajectories are shown, associated with terminal fates: Pericytes, Cardiac Fibroblasts (CFb), and valvular interstitial cells (VICs). Plot sections are coloured to represent distinct phases along the EMT and differentiation trajectory, progressing from Heart Mesothelium/Epicardium to EPDC, and subsequently to terminal fates. Vertical lines indicate the boundary pseudotime values demarcating these trajectory phases. MAGIC imputed expression values plotted. (H) Violin plot of normalised *Krt7* expression in mesothelia, CM, and CECs. (I) Genome tracks of the *Krt7* locus showing a putative enhancer (highlighted). (j) Normalised ATAC-seq signals of the CRE from (I) in FACS sorted epicardial cells and EPDCs. Positions of TCF21 binding sites in the enhancer are shown. (K) Imputed expression of *Krt7* along the epicardial EMT and differentiation pseudotime trajectory. (L) Violin plot of normalised *Upk3b* expression in mesothelia, CM, and CECs. (M) Genome tracks of the *Upk3b* locus showing a putative enhancer and promoter (highlighted). (N) Normalised ATAC-seq signals of the CRE from (M) in FACS sorted epicardial cells and EPDCs. Positions of TCF21 binding sites in the enhancer are shown. (O) Normalised ATAC-seq signals of the promoter from (M) in FACS sorted epicardial cells and EPDCs. The WT1 motif is shown. (P) Imputed expression of *Upk3b* along the epicardial EMT and differentiation pseudotime trajectory. *CM: ventricular cardiomyocytes, EC: endothelial cells, CFb: Cardiac fibroblast, EPDC: epicardium-derived cell, Heart meso: Heart mesothelium, VIC: Valvular interstitial cell* **See also Figure S8**

We also documented conserved regulation of *Krt7*, potentially via a TCF21-linked enhancer (Figure 6H, 6I) that is less accessible in EPDCs (Log2FC: −2.30, FDR: 7.68*10-7; Figure 6J), mirroring *Krt7*’s downregulation during EMT (Figure 6K). While TCF21 could regulate this CRE in the heart mesothelium, its limited expression elsewhere (Figure S8A) suggests that other TFs and CREs are involved. The regulation of *Upk3b*, a robust mesothelial marker (Figure 6L), exemplifies this tissue-specific control. In the epicardium, *Upk3b* is likely regulated by an enhancer linked to the TF TCF21 (Figure 6M, 6N). In other tissues, however, its expression may be controlled by different mechanisms, such as other TFs acting on the same enhancer, or by the pan-mesothelial TF WT1 binding directly to the *Upk3b* promoter (Figure 6O). Both *Upk3b*-linked elements showed significantly decreased accessibility in EPDCs (Figure 6N, 6O), correlating with *Upk3b*’s downregulation during EMT (Figure 6P).

These CREs also appear to drive temporal expression. *Maf* and its target genes *Krt8* and *Krt18* all have increased expression as the epicardium matures (Figure S8B), as does the accessibility of their respective CREs (Figure S8C, S8D). On the other hand, *Upk3b* expression shows limited temporal variation (Figure S8B). This finding is consistent with the accessibility of *Upk3b*-linked elements (Figure S8E), closely mirroring expression and reinforcing its regulatory role. In the pancreas, the *Krt18*-linked enhancer’s accessibility mirrors the gene’s complex expression pattern (downregulation by E14.5 followed by upregulation by E17.5), and *Maf* expression generally correlates with *Krt8* and *Krt18* (Figure S8F, S8G, S8H). For *Krt8*, a contribution from other regulatory elements, such as the *Krt8* promoter, is substantial, as *Krt8* promoter accessibility temporally correlated more strongly with gene expression (Figure S8H). Unlike in the epicardium, *Upk3b* expression in the pancreas mesothelium increases over time (Figure S8F), which correlates with the accessibility values of both *Upk3b*-linked elements (Figure S8I). Similar temporal correlations between CRE accessibility and gene expression for *Krt8*, *Krt18*, *Maf*, and *Upk3b* were also observed in the lung (Figure S8L-O). Overall, our data suggest MAF governs cytokeratin expression and mesothelial identity in multiple mesothelia.

### Cross-species comparison supports the role of MAF and enables a re-evaluation of epicardial markers

To test if these regulators, which may modulate gene expression in both epicardium-specific and pan-mesothelial manners in mouse, play similar roles in humans, through multiple rounds of single-cell integration (Figure 7A), we generated a cross-species integrated dataset of mouse (E10.5-P6) and human (PCW 6-22) epicardial cells. After benchmarking integration methods (Supplemental Table 12), we used scVI ^76^ to harmonize the datasets, yielding an integrated scRNA-seq dataset of 5,784 epicardial cells (Figures 7B, S9A, S9B).

**Figure 7.**
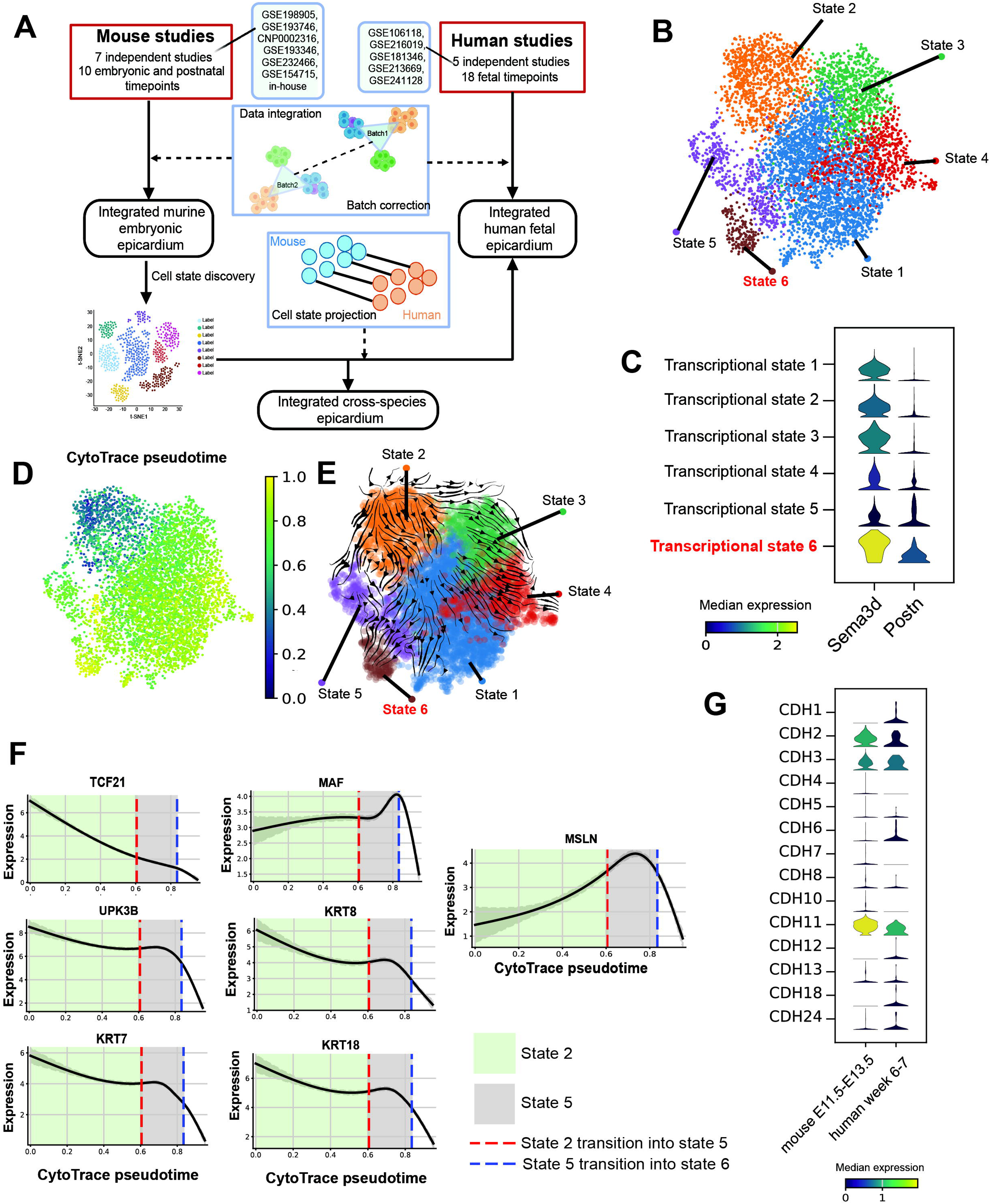
Human and mouse cross-species comparison of transcriptional regulators and gene markers in the epicardium. (A) Schematic of the cross-species comparison methodology. (B) UMAP of the integrated human-mouse embryonic/fetal epicardial cells, showing transcriptional cell states. (C) Stacked violin plot of median SEMA3D and POSTN expression across cell states. (D) UMAP showing CytoTrace pseudotime of epicardial cells. (E) CellRank transition matrix UMAP showing predicted trajectories between cell states. (F) Imputed expression of MAF, TCF21, and their target genes along the inferred epicardial EMT pseudotime trajectory. A trajectory associated with epicardial cells initiating EMT and losing mesothelial features is shown. Plot sections are coloured to represent distinct phases along the trajectory, progressing from State 2 to State 5 and State 6. Vertical lines indicate the boundary pseudotime values demarcating these trajectory phases. MAGIC imputed expression values plotted. (G) Stacked violin plots of median cadherin expression in mouse (E11.5-E13.5) and human (PCW 6-7) epicardial cells. **See also Figure S9**

Clustering, gene marker identification, and GO pathway enrichment identified states for non-proliferative (State 1), proliferative (State 2; marked by MKI67), lubrication (State 3, marked by PRG4 and MUC16), and epicardial adipose tissue (State 4, marked by ITLN1) (Figure S9C, S9D). We also captured a migratory cell state (State 6), similar to one described in a human fetal heart atlas ^77^, which highly expresses POSTN and SEMA3D, indicative of migration and paracrine control of cardiac innervation, roles associated with EPDCs ^78–80^ (Figure 7C). This state is likely resolvable due to the multi-layered nature of the human ventricular epicardium ^49, 50^. We hypothesise this structural difference enables the resolution of very early EMT stages in human tissue, as cells retain some epicardial characteristics during their prolonged migration. In mouse, the rapid transition from epicardium to sub-epicardium likely obscures these initial phenotypic changes.

Using CytoTrace ^81^ to infer developmental potential (Figure 7D), we uncovered a trajectory from proliferating epicardium to an intermediate state (State 5) and then to the migratory subpopulation (Figure 7E). Along this trajectory, MAF, TCF21, and their targets (KRT8, KRT18, *KRT7*, and UPK3B) were downregulated as cells lost their mesothelial identity, evidenced by the loss of MSLN expression, and supporting their roles in maintaining mesothelial identity (Figure 7F).

The cross-species dataset confirmed that our inferences on cadherins are conserved. Analyzing mouse epicardial cells from E11.5 to E13.5, and human cells at approximately equivalent developmental stages (post-conception week 6 to 7), CDH2, CDH3, and CDH11 are expressed in both human and mouse epicardium (Figure 7G). Importantly, E-cadherin/CDH1 shows species-specific expression: it is expressed in human epicardial cells (albeit at a low level compared to other cadherins) but is absent in the mouse epicardium from E10.5 to P6 (Figure 7G, S9E). Using our integrated murine heart atlas (Figure S9F), we confirmed *Cdh1* expression is absent from the entire mouse heart (Figure S9G), consistent with Figure S1H and in line with another mouse heart atlas ^82^ (Figure S9H). CDH1 is likely silenced by H3K27me3 modification near its promoter (Figure S9I), a process potentially driven by PRC2 binding. PRC2 binding to the *Cdh1* promoter has been demonstrated in mouse E14.5 ventricular apex ^83^. PRC2-mediated heterochromatin deposition may contribute to cross-species variation, offering an avenue to improve the translatability of non-human models. Altogether, our cross-species dataset verified the human relevance of our findings and represents a valuable resource for cardiac biology.

## Discussion

The coelomic mesothelium’s role in visceral morphogenesis underscores its potential for organ regeneration. Mesothelial dysfunction contributes to congenital diseases ^77, 84–87^, fibrosis ^88–90^, cardiomyopathies ^91^, and mesothelioma ^92, 93^. Despite its importance, the gene regulatory logic governing mesothelial identity remains elusive. Decoding these GRNs would improve understanding of this layer’s therapeutic importance and provide a basis for reactivating the quiescent adult mesothelium for regeneration ^94^. We present a cross-organ comparison to examine conserved and organ-specific mesothelial roles. We find *Upk3b* robustly marks the mesothelial compartment, suggesting it could be used for lineage tracing to overcome the non-specific expression of *Wt1* in coronary endothelial cells ^95^. We further inferred mechanisms behind *Upk3b* regulation mediated by a series of CREs and TFs, including WT1.

Our analysis shows each organ’s mesothelium has a distinct gene expression and regulation landscape. We identified enriched markers that distinguish mesothelia from each other and from other cell types in their organ (e.g., *Pnoc* for lung, *Pitx2* and *Barx1* for pancreas). Our initial finding of *Msln* being exclusive to the E13.5 epicardium reflects the earlier morphogenesis of the heart; the gene is expressed later in lung and pancreas mesothelia. We propose *Msln*’s spatiotemporal expression is regulated by a conserved set of CREs, potentially driven by common TFs such as WT1 and NRF2.

Our investigation of the lung mesothelium revealed new regulatory complexity. We identified FOXF1 as a potential upstream regulator of the Hedgehog signal transducer *GLI1*, linking it to lung development. This is supported by similar phenotypes (lung hypoplasia) from *Foxf1* haploinsufficiency, mesenchyme-specific *Foxf1* inactivation, and disruptions to Hedgehog signaling ^37, 96, 97^. These phenotypes may result from disrupted mesothelial EMT. As FOXF1 is also a known Hedgehog target, our findings suggest a potential bidirectional interplay between FOXF1 and the Hedgehog signaling pathway.

Our analysis also unveiled new insights into the developing pancreas’s genomic landscape. We propose TFs such as PITX2, ISL1, and BARX1 mediate mesothelial function via CREs whose activity correlates with temporal gene expression. In particular, PITX2 may be critical for cell layer integrity via *Cdh6* and may influence exocrine pancreas development through its proposed regulation of *Igf1* ^57^. BARX1’s potential regulation of *Hoxb8* may have implications for pancreas organogenesis and warrants further exploration, as other HOX family members such as HOXB6 are involved in endocrine cell differentiation ^98^.

A comparative analysis also identified key factors contributing to epicardial function and identity. We identified the TBX20 regulon, which may control genes with critical roles in the embryonic epicardium, including *Csf1, Sema3d, Fgf2*, and *Rarres2*. Through these genes, TBX20 may influence epicardial behaviours such as establishing immune niches, axon pathfinding, and EMT, which should be investigated further. We also proposed that TBX20 regulates plakophilin-2 (*Pkp2*), a key desmosomal component that protects tissues under mechanical stress and may safeguard epicardial identity. *Pkp2* expression may therefore negatively correlate with EMT, as its loss can increase cell migration ^99^.

Besides desmosomes, epicardial cells rely on adherens junctions formed by cadherins like *Cdh2, Cdh3*, and *Cdh11* to maintain intercellular junctions and layer integrity. Through an integrated, cross-species analysis, we proposed a negative role for *Cdh3* and *Cdh11* in EMT and confirmed their expression in the human epicardium. Our *in silico* approach also clarified the contested role of *Cdh1* (E-cadherin). While present in the human epicardium at low levels, its transcription in the murine heart is negligible, likely due to epigenetic silencing. Using CDH1 as the case study, we suggest that divergent gene expression patterns across species may arise due to adaptive evolutionary genomics. Nevertheless, our analysis highlights the cross-species similarity between the mouse and human embryonic epicardium, evident by transcriptomic state conservation and similar cadherin expression.

Mesothelial biology may be conserved not only across species but also organs. By investigating the conserved regulatory architecture of the coelomic mesothelium, we identified MAF as a potential gatekeeper of mesothelial identity across organs. We showed a negative association between MAF and EMT in both the mouse and human epicardium. We then confirmed the conserved nature of mesothelial enhancers that may be regulated by MAF. Through these enhancers, MAF may modulate cytokeratin genes such as *Krt8* and *Krt18*, which form part of the keratin cytoskeleton connected to desmosomes ^100^. This finding is consistent with previous observations that identify MAF as a driver of epidermal progenitor differentiation into keratinocytes by regulating keratin gene expression ^101^.

Overall, our work provides the first comprehensive overview of the gene regulatory landscape of the coelomic mesothelium. By leveraging cell-type-specific and cross-species genomics, we provide initial insights into organ-specific and conserved mesothelial regulatory programs. We identify high-confidence enhancers that can be manipulated to gain a deeper understanding of mesothelial biology, laying the groundwork for future studies on the mesothelium’s role in disease and regeneration.

### Limitations of the study

Our study relies on computational predictions of E-P connections from transcriptomic and epigenomic data, which require experimental validation. Data availability was a limitation, particularly the lack of chromatin conformation data (e.g., Hi-C) for mesothelial cells to confirm E-P contacts. Our histone CUT&RUN-seq data were from cultured epicardial cells, which may differ from the *in vivo* epigenome. While we identified plausible enhancers for each gene, other regulatory elements may contribute to gene expression. Our cross-species inferences were limited by the current lack of human epicardial ATAC-seq data and limited RNA-seq data from early, EMT-relevant developmental stages. A more comprehensive cross-species comparison will require techniques such as Hi-C and cell-type-specific mapping of PRC2 binding.

## Acknowledgements

The Dunn School and Ludwig Institute for Cancer Research Flow Cytometry facility for advice and assistance, and Biomedical Services staff for animal husbandry. This work was funded by the British Heart Foundation (BHF) Intermediate Basic Science Research Fellowship (JMV: FS/19/31/34158), BHF Ian Fleming Senior Basic Science Research Fellowship (NS: FS/19/32/34376) and Oxford BHF Centre of Research Excellence awards to JMV and ANR, and to NS and ANR (both RE/18/3/34214). QD is funded by a doctoral training scholarship from Vinmec International Healthcare System.

## Author Contributions

Conceptualisation, QD, ANR, JMV; Methodology, QD, ANR; Investigation, QD, ANR; Formal Analysis, QD, ANR; Visualisation, QD; Writing – Original Draft, QD; Writing – Review and Editing, QD, ANR, NS, JNV; Funding Acquisition, ANR, NS, JNV; Supervision, ANR, JNV, NS.

## Declaration of interests

The authors declare no competing interests.

## STAR Methods

### Key resources table

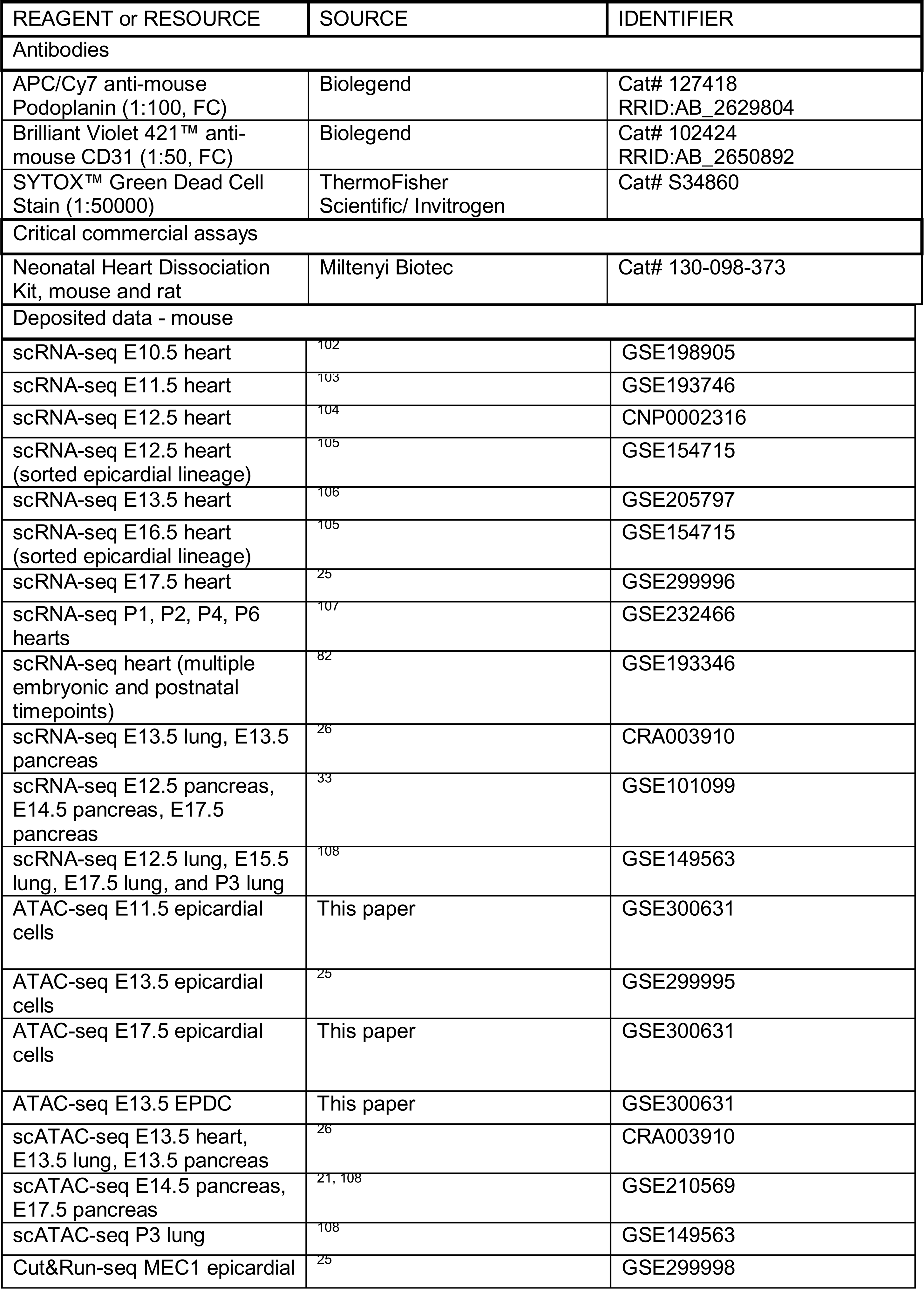
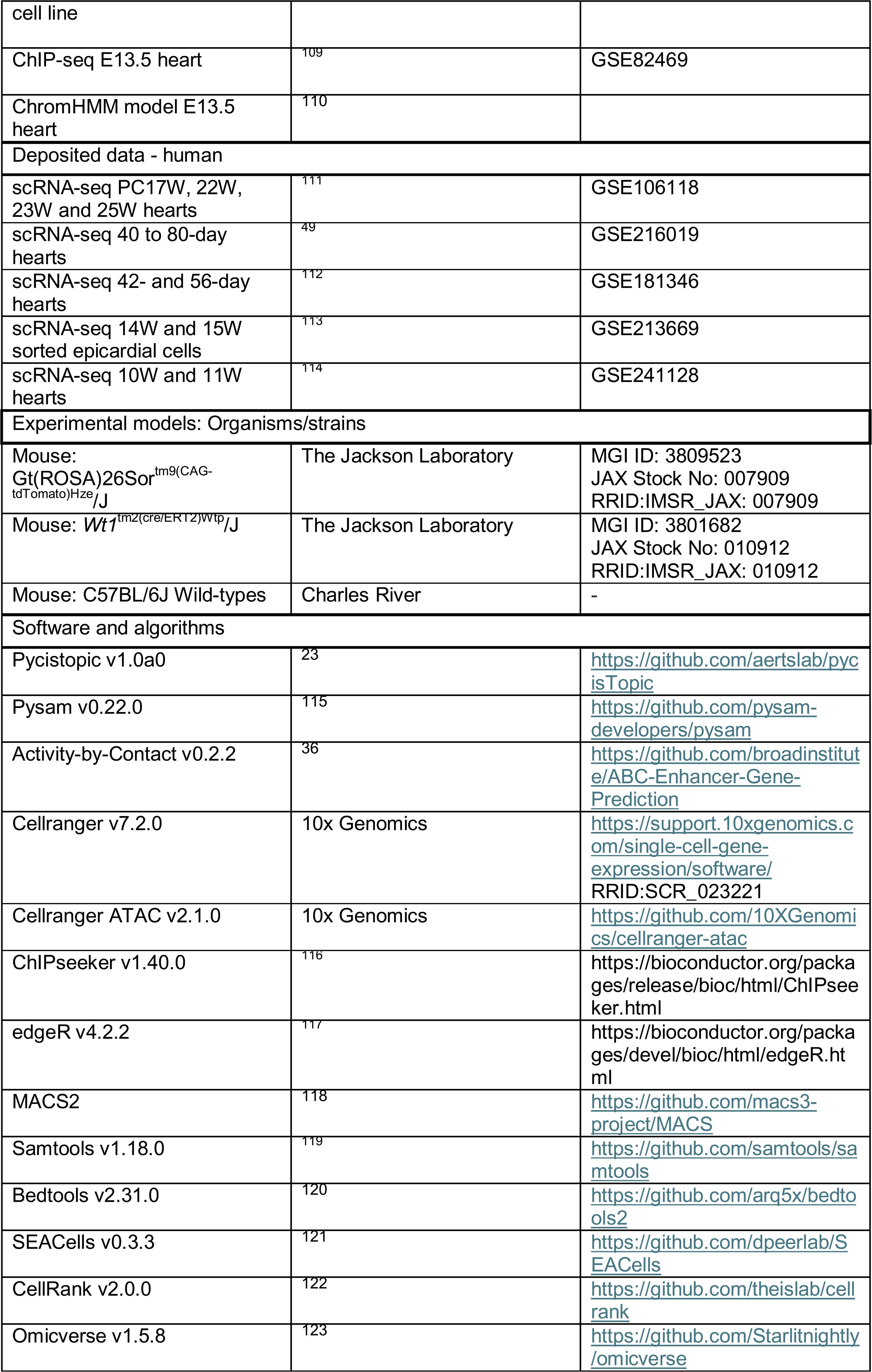
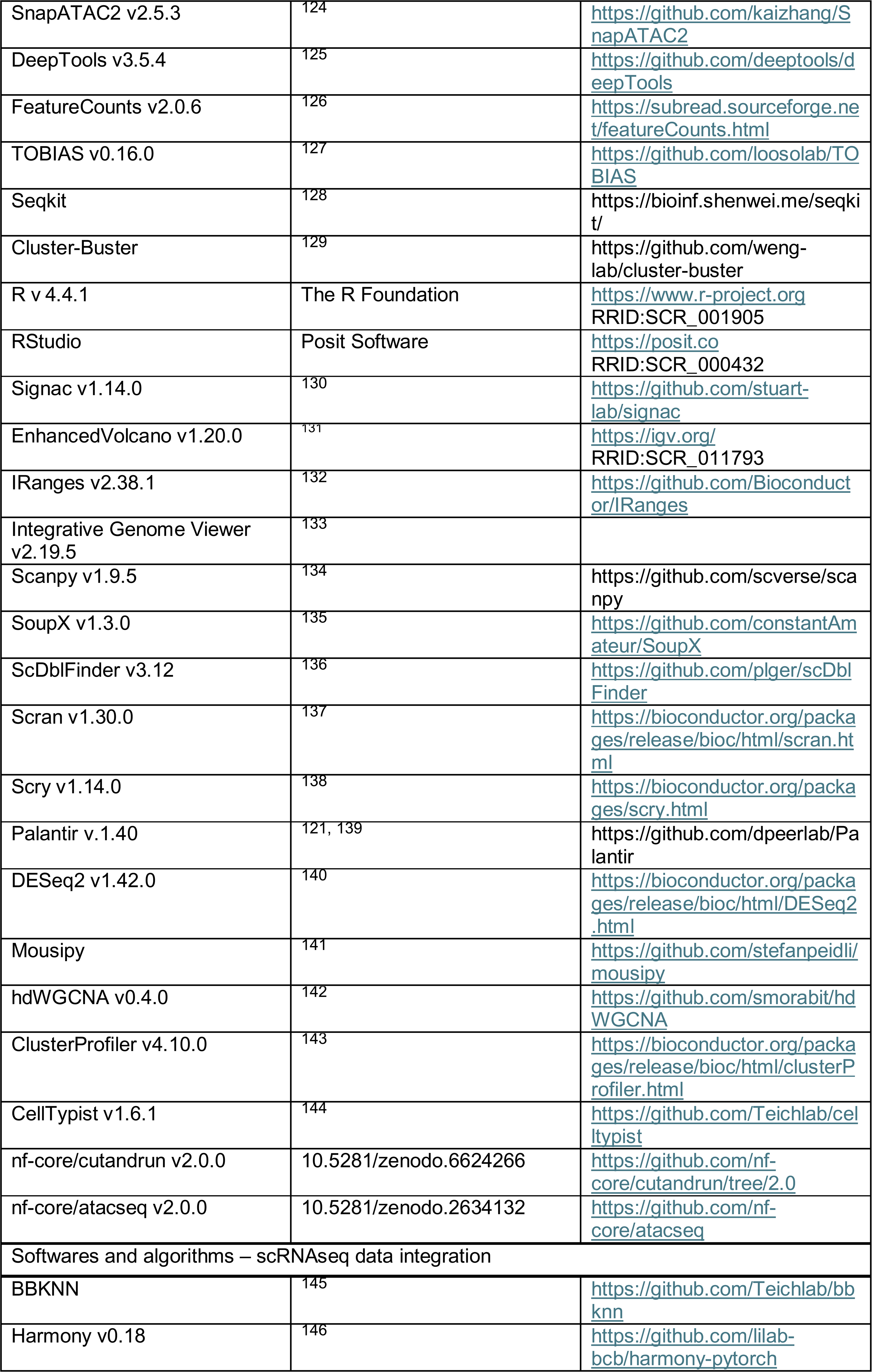
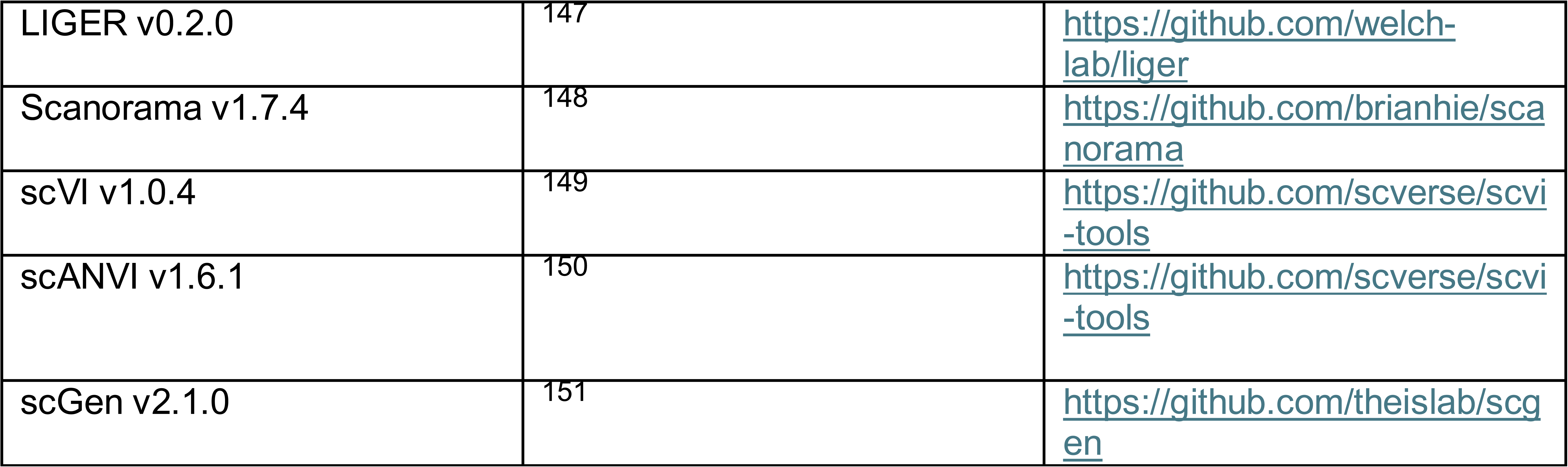

### Resource availability

#### Lead contact

Further information and requests for resources should be directed to the lead contact, Joaquim M. Vieira.

#### Data and code availability

ATAC-seq data have been deposited at GEO as GSE300631 and are publicly available as of the date of publication.

This paper analyzes existing, publicly available data, as detailed in STAR Methods.

Original code and custom pipelines have been deposited at https://github.com/loganminhdang/Mesothelium_paper_2025.

Any additional information required to reanalyse the data reported in this work paper is available from the lead contact upon request.

### Method details

#### Animals

##### Mouse Strains

Males homozygous for *Gt(ROSA)26Sortm14(CAG-tdTomato)Hze/J* or *Gt(ROSA)26Sortm9(CAG-tdTomato)Hze/J* (*Rosa26tdTomato*), and heterozygous for *Wt1tm2(cre/ERT2)Wtp/J* (*Wt1^CreERT2/^*^+^) were crossed with *Rosa26tdTomato* or C57BL/6J wild-type females for genetic-lineage tracing studies. Pregnant females were administered 100 mg/kg tamoxifen by oral gavage at E9.5. All procedures were approved by the University of Oxford Animal Welfare and Ethical Review Board, in accordance with Animals (Scientific Procedures) Act 1986 (Home Office, UK).

##### Fluorescence-activated cell sorting (FACS)

E11.5, E12.5 or E13.5, and E17.5 hearts (*Wt1^CreERT2/+^;Rosa26tdTomato*) from littermate embryos were pooled prior to dissociation (E11.5: 6 hearts/sample; E13.5: 9-13 hearts/sample; E17.5: 4 hearts/sample) to maximise cell numbers for FACS and corresponding FMO controls, and achieve a target sort of 50,000 epicardial cells/sample for the downstream ATAC-seq protocol. Single-cell suspensions in cell staining buffer (Biolegend) were pre-treated for 5 minutes on ice with TruStain FcX antibody, followed by surface staining with BV421 anti-CD31 and APC/Cy7 anti-Podoplanin (PDPN) antibody for 30 minutes on ice, then washed with PBS before being incubated with SYTOX Green Dead Cell Stain for 5 minutes on ice. After PBS washes, cells were resuspended in sorting buffer containing 2% FBS in PBS. Primary antibodies and SYTOX stain used are detailed in the Key Resources Table. BD FACSAria III was used to sort 10,000-50,000 live cells negative for SYTOX Green and CD31, and positive for tdTomato and PDPN, into 0.5% BSA in PBS.

##### ATAC-seq data generation

Using FACS, we isolated live epicardial cells (CD31-tdTomato+PDPN+) from E11.5 and E17.5 hearts, and EPDCs (CD31-tdTomato+PDPN-) from E12.5 and E13.5 hearts (E12.5 and E13.5 hearts were both used to represent the E13.5 stage due to very strong concordance between these two stages). Cells were subjected to the ATAC-seq protocol as previously described ^152^. In brief, 10,000-50,000 sorted cells were centrifuged at 500 ×g. for 5 minutes, the supernatant was removed, and the pellet was resuspended in 50µL of ATAC Resuspension Buffer (RSB: 10mM Tris-HCl pH 7.5, 10mM NaCl, 3mM MgCl2) supplemented with 0.1% NP-40, 0.1% Tween-20, and 0.01% Digitonin to extract nuclei. After a 3-minute incubation, 1mL of ATAC RSB with 0.1% Tween-20 was added, mixed by inversion, and centrifuged at 500 ×g for 10 minutes. The supernatant was carefully removed, and nuclei were resuspended in 50µL of transposition mix (100mM tagmentation enzyme, 0.01% Digitonin, 0.1% Tween-20, 0.33X PBS, in TD buffer). The tagmentation enzyme amount was adjusted according to cell count (for 40,000 cells, the enzyme was diluted to a 0.8× concentration in TD buffer before being added to the transposition mix). Incubation and centrifugation steps were performed on ice or at 4°C, respectively, up to this point. Transposition reactions were carried out at 37°C for 30 minutes with shaking at 1,000 RPM using an Eppendorf ThermoMixer.

ATAC-seq DNA libraries were generated as previously described ^152^, with minor modifications. Reactions were purified using the MinElute Reaction Cleanup Kit (Qiagen) following the manufacturer’s instructions. Pre-amplification of transposed fragments (5 cycles) was performed using NEBNext Ultra II Q5 Master Mix (NEB) and Nextera i7 index adapters (IDT). The number of additional amplification cycles (typically 4) was determined using the qPCR method ^153^. Final PCR products were purified using a double-sided bead purification method with SPRIselect beads (0.5X volume followed by 1.3X volume of SPRI beads). Final cleanup was performed with the MinElute Reaction Cleanup Kit (Qiagen). Libraries were assessed for quality using an Agilent TapeStation with a D1000 DNA ScreenTape assay, and quantified with the KAPA Library Quantification Kit. Next-generation sequencing (NGS) was conducted on equimolar pooled libraries using the NextSeq 500/550 High Output v2.5 kit (75 cycles) (Illumina), with paired-end reads (40 bp x 2) and 8 bp single index reads.

##### Bulk ATAC-seq data processing

Briefly, the ATAC-seq nf-core Nextflow pipeline (version 2.0) ^154^ was used to pre-process ATAC-seq sequencing data, consisting of the following steps: quality control, adapter trimming, alignment to the UCSC mm10 genome, filtering unwanted reads, and merging across biological replicates. Filtering removed: reads aligning to mitochondrial DNA and blacklisted regions ^155^: PCR duplicates, multi-mapped reads, and reads that have an insert size > 2kb and contain > 4 mismatches.

Alignment files were converted to bed files using the randsample function of MACS2 (version 2.2.9.1) ^118^ with 100% of tags kept (-p 100). Peak-calling was performed with MACS2 using the following parameters: -f BEDPE -g mm -q 0.01 --nomodel --shift −73 --extsize 146 --call-summits --cutoff-analysis --keep-dup all.

The resulting peak summits were then extended by 250 bp to both sides to a fixed width of 501 bp. Peaks that extend beyond chromosome ends were filtered. The remaining peaks went through an iterative removal process to filter out less significant peaks that overlap with more significant ones, as described in ^30^. For joint analysis of all developmental stages, we merged biological replicates for each stage into a single bam file represent each stage, then sort and index using Samtools (version 1.18.0) ^119^. We then merged the sorted bam files of E11.5, E13.5, and E17.5 samples into a single bam file using the samtools merge function. MACS2 peak calling and processing were performed exactly as the procedure described above. Intersection of peaks between samples was performed using either the bedtools intersect function of Bedtools (version 2.31.0) ^120^ or the findOverlaps function of the IRanges library (version 2.38.1) ^132^.

##### Single-cell RNA-seq data pre-processing

Single-cell RNA-seq datasets were processed via CellRanger (version 7.2.0) using the mm10 reference and underwent quality control and normalisation according to published best practices ^156^. Briefly, we filtered out low-quality cells, which are defined as barcodes that: (1) are outside the range of 5 median absolute deviations (MADs) for count depth and genes per barcode; (2) outside of 3 MADs for the fraction of counts from mitochondrial genes per barcode; (3) cells with a percentage of mitochondrial counts exceeding a manual selected threshold. We used a dynamic threshold for percentage of mitochondrial counts due to the observed large proportion of mitochondrial DNA in cardiac cell types ^157^, ranging from 0 to 8%. For non-cardiac datasets, we used a threshold of 15%. Next, ambient RNA correction and doublet detection were done via soupX (version 1.3.0) ^135^ and scDblFinder (version 3.12) ^136^, respectively. Single-cell counts were normalised using the shifted logarithm approach implemented in Scanpy (version 1.9.5) ^134^. To select highly variable genes (HVG), for individual datasets, corrected counts were normalized using the Scran’s pooling-based size factor estimation method implemented in the scran (version 1.30.0) R library ^137^. Scran-normalized counts were then used to select top 4000 HVGs via the R scry (version 1.14.0) library ^138^. Dimensionality reduction and clustering were performed using the UMAP ^158^ and Leiden ^159^ algorithms, respectively. We calculated the UMAP embedding by finding nearest neighbors in the principal component analysis space.

##### Single-cell ATAC-seq data processing

Raw sequencing files were processed via the CellRanger-ATAC pipeline (version 2.1.0) using the mm10 reference. Pre-processing was performed using SnapATAC2 (version 2.5.3) ^124^. Briefly, a minimum TSS enrichment of 10 and a minimum number of fragments of 4000 was chosen to filter for high-quality cells. After removing features that map to blacklisted regions, we count fragments instead of reads to better preserve regulatory information, as demonstrated in Martens et al. (2024) ^160^. Doublet removal was performed prior to dimensionality reduction via spectral embedding and clustering via the Leiden algorithm.

For peak calling, ATAC-seq fragments for each cell type were pseudo-bulked using the function export_pseudobulk of the Pycistopic package (version 1.0a0), based on the UCSC mm10 genome. Peak-calling was performed on the pseudo-bulk profiles, using the following parameters: --format BEDPE --gsize mm --qvalue 0.01 --nomodel --shift 73 --extsize 146 --keep-dup all --call-summits –nolambda. Consensus peaks were inferred using the summit extension and iterative filtering approach used in bulk ATAC-seq data processing.

##### ATAC-seq data analysis

To generate a list of consensus non-overlapping peaks across tissue mesothelia, the procedure from Corces et al. (2018) ^30^ was implemented. Briefly, to normalize the differences between samples in terms of read depth and quality, the individual MACS2 501-bp peak score (“-log10(pvalue)”) was divided by the sum of all peak scores in each sample and multiplied by 1 million. The peak sets from tissue mesothelia were then merged using Bedtools and iteratively filtered again to retain significant peaks from all samples to generate a master peak set representing producible mesothelial peaks.

BigWig files used to calculate the correlation between organ mesothelia and cardiac cell types were generated using Pycistopic. Average scores for consensus mesothelial peaks between bigWig files were calculated using the multiBigwigSummary function of DeepTools (version 3.5.4). Overall similarity between samples was determined by computing Pearson correlation coefficients based on read coverage in consensus peaks. Sample signals over the set of consensus peaks were calculated using the computeMatrix function of Deeptools ^125^ using the following parameters: --referencePoint center -b 500 -a 500 – missingDataAsZero and visualised using the plotHeatmap function.

The bigWig files for the epicardium bulk ATAC-seq, including the E13.5 stage, were normalised using the trimmed mean of M values normalisation method to account for differences in the peak landscape (signal-to-noise ratio) between samples. We used FeatureCounts to extract read counts for peaks and create a region-by-sample matrix. We then used this matrix to calculate a scale factor for each sample, which was then used to scale bigWig files accordingly using the –scaleFactor parameter of the bamCoverage tool.

##### Differential chromatin accessibility analysis

We adapted the procedure from Reske et al. (2020) ^161^ and conducted differential analysis using three biological replicates for each sample (epicardium: E11.5, E13.5, E17.5; EPDC: E13.5). To select for peaks and normalisation method, we tested four peaking calling and normalisation strategies as recommended by the original report (MACS2 + TMM normalisation, MACS2 + loess normalisation, csaw *de novo* peak identification by local enrichment + TMM normalisation, and csaw *de novo* peaks + loess normalisation). For MACS2, we used peak-calling parameters and iterative peak filtering strategy as described in the “ATAC-seq data analysis” method section. For identifying *de* novo peaks using csaw, we performed local enrichment by counting BAM reads in 300 bp window and filtering features by local enrichment (threshold of 3-fold increase in enrichment over 2kb window). We found that our MACS2-dependent peak-calling strategy provides comparable performance to the csaw peak identification. TMM normalisation, when used in conjunction with our MACS2 strategy, does not create global normalisation bias.

For differential accessibility analysis, we used edgeR (version 4.2.2) ^117^. Using the ATAC-seq count matrices with TMM-normalised factors, dispersions were stabilised by empirical Bayes, followed by quasi-likelihood GLM fitting for hypothesis testing for differentially accessible windows (500bp). Windows were merged, with the most significant windows retained as the statistical representation. FDR threshold for detecting significantly differentially accessible peaks was set at 0.05. Differential peaks were annotated with gene information using the ChIPseeker R library (1.40.0) ^116^, using the UCSC mm10 database.

##### Single-cell RNA-seq batch correction and integration

For batch correction and data integration of multiple datasets (mouse embryonic epicardium, mouse embryonic and postnatal lung mesothelium, mouse embryonic pancreas mesothelium), we downsampled large datasets to a maximum of 10,000 cells using the “subsample” function in Scanpy. HVGs were selected in a batch-aware manner using Scanpy by calculate HVGs for each batch separately and select HVGs that are highly variable in the highest number of batches. Using only HVGs, we ran seven selected data integration methods according to default parameters obtained from paper methods and tutorials. For further details on the methods, please see Supplementary Table 12. As some methods, such as scANVI or scGen, required prior cell type annotation, we annotated cells in the concatenated dataset prior to integration. For cardiac samples, we used automated annotation via CellTypist (version 1.6.1) ^144^ based on cell-type annotation from existing datasets ^82^ for mouse and ^112^ for human). Briefly, we normalised raw counts from these datasets to 10000 counts per cell, then transformed via shifted logarithm. Via Celltypist, a logistic regression model was trained using filtered genes as features. We then used this model to annotate cell types in single-cell datasets, followed by careful manual inspection of marker genes to ensure correct annotation. For other datasets, cell type annotation was performed manually by matching cluster-specific genes to established transcriptomic markers.

We performed benchmarking of integration based on the procedure described in Luecken et al. (2022) ^162^, using the metrics grouped into two broad categories: (1) removal of batch effects and (2) conservation of biological variance. For cross-species integration, as we focused exclusively on the epicardium, we ranked methods based on their ability to correct batch effects.

##### Metacell-based differential expression analysis

To circumvent issues associated with single-cell pseudo-bulking approaches, we created metacells – artificial aggregates of single cells. The use of metacells to increase statistical rigor has been applied previously ^163^. In our case, we generated metacells to create statistical dispersion that enhances the performance of differential analysis frameworks such as DESeq2 ^140^.

We used SEACells (version 0.3.3) ^121^ to build transcriptomic metacells for all organ mesothelia. The input is an AnnData object comprising of the entire pre-filtered gene, cell type assignment, and a low-dimensional UMAP representation. To build metacells – coarse aggregate of phenotypically similar cells, we first sampled 10 cells to uniformly cover the phenotypic landscape. We define one metacell per 75 single cells and used a convergence percentage of 1e-5. Following the construction of metacells, we sum the unnormalized soupX-corrected counts of individual cell in each metacell to generate counts of all genes in each metacell. Each metacell was then assigned a cell type label based on which cell type was most prominent amongst individual cell used to construct that metacell. Following the generation of a counts x gene matrix, we treat each metacell as a biological “replicate” and performed differential gene expression analysis using DESeq2 (version 1.42.0). We filtered genes with low count (sum of expression count across all metacells less than 50). Count data was normalized using DESeq2’s built-in median-of-ratios method. Differentially expressed genes between organ mesothelia are considered genes with Bonferroni-Hochberg-adjusted p-value less than 0.05 (to account for multiple comparisons) and Log2FC > 1.

##### Mouse epicardial lineage integration

To mitigate the confounding influence of inter-platform technical variation on biological inference, we selected only datasets generated with UMI-based technologies (10X Genomics). Datasets from other technologies, such as the full-length Smart-seq2 method, which is known to exhibit higher gene detection sensitivity, are deliberately excluded. This intrinsic difference in transcript recovery between technologies can introduce non-biological variance upon single-cell integration, thereby confounding downstream analyses such as pseudotime inference.

We performed quality control on individual datasets, which included ambient RNA correction and doublet detection, before concatenating the datasets and following the steps outlined in the section on batch correction and integration methods. We selected scANVI due to its performance in integration metrics. We extracted the latent representation of each cell generated by scANVI and calculated a new UMAP embedding using this corrected representation. Following a round of clustering using Leiden (resolution 0.5), we identified cell-type marker genes using the Wilcoxon rank-sum test implemented in Scanpy. Cell types associated with epicardial EMT, based on marker annotation and UMAP distance, are then clustered out to form a new subsetted dataset. We identified cell types based on established markers: *Upk3b* (epicardium), *Postn* (EPDC), *Rgs5* (Pericytes), *Dpt* and *Col15a1* (CFb/Fb progenitors), *Adamts19*, *Col9a3*, and *Lef1* (VIC), and *Myl9* (SMC progenitor) ^164–169^.

Further data cleaning was performed to ensure epicardial lineage by only retaining cells with tdTomato reporter expression. Gene expression counts were then re-normalised using the shifted logarithm method. A further round of marker gene identification was then performed to identify cell states.

##### Cross-species data integration

For each species, we pre-processed individual datasets as outlined earlier and integrated them using the best-performing method according to our benchmarking (scGen for mouse embryonic/postnatal heart and scANVI for human fetal heart). Following integration, we clustered out epicardial cells based on their expression of WT1 and UPK3B. For the human fetal heart, as the epicardial cluster is confounded by the presence of cells expressing high levels of haemoglobin genes, we performed multiple rounds of clustering to remove these cells and minimise noise in the dataset. We then concatenated the clustered-out mouse and human epicardial cells into a single dataset. To harmonise human and mouse gene symbols, we mapped mouse (GRCm39) genes to their corresponding human (GRCh38.p13) orthologs using mousipy ^141^. We then normalised raw counts in the combined dataset using the shifted log approach, followed by dimensionality reduction and clustering. 5000 HVGs were selected on the basis of being highly variable in at least one species. Data integration was then performed using HVGs. We only selected methods that are agnostic to cell-type label and found scVI outperformed at correcting species difference (using raw counts, a 30-dimensional latent space and two hidden layers for encoder and decoder neural networks). We used the cell latent representation generated by the scVI model to compute a new UMAP embedding.

##### Gene co-expression network inference

We applied hdWGCNA ^142^ to scRNA-seq dataset consisting of three organ mesothelia to detect network modules of genes whose expression profiles are tightly intertwined. Briefly, we construct metacells based on the low-dimensional embedding of the dataset (aggregating 75 cells for each metacell). We then constructed a gene-gene correlation adjacency matrix to infer co-expression relationships between genes, selecting a soft power threshold of 5 to reduce noise and retain only strong correlations. Modules were projected from the murine mesothelial scRNA-seq reference to murine mesothelial ATAC-seq query using the hdWGCNA function ProjectModules. We assessed module preservation using the Z statistic ^32^. Over-representation analyses based on regulon target genes were done via the enrichGO function of R library ClusterProfiler (version 4.10.0) ^143^, using the GO Biological Process database and with Benjamini-Hochberg (BH)-adjusted p-value and q-value cutoff of 0.01 and 0.05 respectively.

##### Differentiation potential analysis

For the integrated mouse epicardial lineage dataset, we used Palantir (version 1.4.0) ^139^. 20 diffusion components were computed to determine the diffusion map of the data, followed by low-dimensional embedding and imputation. We pre-selected cells at the extreme for diffusion components as initial and terminal cells before computing pseudotime.

For inferring cellular progression across the integrated cross-species dataset, we utilized CytoTRACE ^81^. This choice was predicated on CytoTRACE’s core mechanism, which primarily leverages transcriptional diversity (i.e., the number of expressed genes per cell) to estimate differentiation potential. We posited that this fundamental cellular property is likely more robustly conserved across species compared to the specific, detailed gene expression programs and precise manifold topologies upon which algorithms like Palantir depend. Indeed, embryonic epicardium is active and proliferative ^24^, thus may express more genes and satisfy the assumption of CytoTRACE. Furthermore, Palantir’s reliance on constructing a continuous low-dimensional manifold from transcriptional similarity can be confounded by inter-species transcriptional divergence. This divergence is prominent, as specific transcriptional states of epicardial cells are enriched in one species. CytoTRACE’s approach, being less directly dependent on the integrity of a unified manifold, is anticipated to offer more reliable pseudotime inference. Based on CytoTRACE pseudotime, we used CellRank (version 2.0.0) to compute a transition matrix and predict the initial cell state ^122^. This initial state was used to predict terminal states. Visualisation of the transition matrix is done by projecting it into the cross-species integrated UMAP embedding.

##### Biological pathway analysis

We used the AUCell algorithm ^170^ to score the activity of gene sets associated with GO terms (Biological Process 2021 database). Visualisation of epicardial cell state-specific pathways was done using Omicverse (version 1.5.8) ^123^.

##### Enhancer-driven gene regulatory network inference

Cis-regulatory topics were identified using Pycistopic (version 1.0a0). Briefly, topic modelling was performed to infer cis-regulatory topics, which we then used to infer differentially accessible regions and gene activity. In the case of bulk ATAC-seq samples, we simulated single cells to generate sparsity to enhance the performance of topic modelling. Using Pysam (version 0.22.0), single cells were simulated by randomly sampling 20000 reads from the pool of bulk ATAC-seq aligned reads and using 2000 reads as an alignment profile for a single cell. Following topic discovery and annotation (topic threshold 0.2), accessibility of regions was imputed (scale factor 10^6^), which generated a binarised cell-peak matrix as input. From here, differentially accessible regions were identified between organ mesothelia and cardiac cell types using the Wilcoxon rank-sum test.

For motif enrichment to identify TF cistromes, we first generated two cisTarget databases based on two consensus peak sets, one from mesothelial across organs and another for cardiac cell types. We supplemented this database by including 1636 consensus mouse TR motifs based on Yi et al. (2021) ^35^. This expansion creates a more comprehensive set of high-quality murine TF motifs, improving the statistical power for the subsequent motif enrichment and GRN analysis. Briefly, we used the UCSC mm10 reference genome and chromosome size to generate fasta files from consensus regions, adding 1kb of background padding, and subsequently generated cisTarget databases using the “create_cistarget_motif_databases” function.

Combining motif database and processed regions, inference of enhancer-driven Regulons was done using the SCENIC+ method (version 1.0a1) in unpaired multiome mode and “mmusculus” as the species. Pseudogenes and ribosomal genes were excluded from the analysis. We focused exclusively on regulons of the activator archetype, which include TFs that bind to and open the chromatin of target genomic regions to promote the expression of target genes. We did not include repressive elements. Negative associations between TF-CRE-genes do not allow reliable inference of transcriptional repressors, as they can be confounded by the data sparsity inherent to scATAC-seq. High-quality regulons are filtered according to their triplet ranking and metrics comprising this ranking (high importance_R2G and importance_TF2G).

Regulon specificity scores were computed as described previously ^23^. In each regulon, only the top 50 target genes (ranked by triplet score) were considered. Over-representation analyses based on regulon target genes were done via the enrichGO function of R library ClusterProfiler (version 4.10.0).

##### Activity-by-Contact analysis

Epicardial enhancer-promoter connections were predicted using the Activity-by-Contact (ABC) model (version 0.2.2). A custom gene and promoter annotation files were compiled by combining NCBI RefSeq annotation for mm10 genome (build 38.1) with the ENSEMBL mouse database (BiomaRt version 2.58.2). Using NCBI RefSeq gene body annotation, we computed gene length and normalised pseudo-bulked epicardial expression profile to create TPM-normalised epicardial expression. Pseudo-bulked expression for each gene was computed as the sum of expression of each epicardial cell in a sample (i.e., E13.5).

Briefly, to infer E-P connections in the E13.5 epicardium, 501-bp candidate enhancer elements were predicted based on ATAC-seq peaks. Enhancer activities within the 150000 strongest elements were quantified using a combination of ATAC-seq, H3K27ac signal, and TPM-normalized gene expression. ABC scores were then computed by combining region activity and Hi-C contact frequency estimated by a power-law function of genomic distance. The powerlaw exponent was set to be 0.87. To flag genes that are impervious to the effect of distal enhancers, a list of ubiquitously expressed genes across mouse tissues was collected from ^171^. To infer E-P links across all developmental stages, we used the consensus epicardial alignment file (merged from all three time points) and filtered peaks called from this file to run the ABC model. The threshold for ABC score is 0.02, which corresponds to roughly 70% recall in human benchmarks.

##### Transcription factor footprint pre-processing and analysis

Calculation of TF footprint scores was done using the TOBIAS pipeline v0.17.0 ^127^. Briefly, the pipeline performs Tn5 Transposase insertion bias correction and footprinting score calculation from the bias-corrected cut sites. Footprint scores across tissue mesothelia were compared to identify TF differential binding. Only motifs annotated to TFs that are expressed in at least one tissue mesothelium were considered.

To identify genes that may be subjected to promoter-mediated regulation, we selected genes that are differentially enriched in the epicardium, in addition to having bound footprints of relevant TFs within the 500 bp region surrounding their TSS.

To scan for all TF binding sites and associated footprints in the *Msln* promoter, we used the *submerge* function of TOBIAS.

##### Epigenomic data analysis

CUT&RUN-seq data was pre-processed as previously described ^25^. Briefly, H3K27ac and H3K4me3 Cut&Run-seq data of the MEC1 mouse epicardial cell line were pre-processed using the nf-core NextFlow pipeline (version 2.0). The following steps were performed: quality control, and alignment to both target and spike-in genomes with a minimum q-score of 10. Peaks were normalised against IgG controls, scaling background control by a factor of 0.8. Read counts were normalised using Counts Per Million (CPM), and consensus peaks, representing active promoters and enhancers, were identified and merged from at least three biological replicates.

Individual replicates were merged, sorted, and indexed using Samtools. Merged bam files were converted into BigWig files using the bamCoverage tool of Deeptools, normalised using the CPM method, with bin size 10. The signal distribution of histone modification was calculated using the computeMatrix function of Deeptools with the following parameters: --beforeRegionStartLength 1000 --regionBodyLength 5000 --afterRegionStartLength 1000 – skipZeros (H3K4me3, active promoters) and --beforeRegionStartLength 3000 --regionBodyLength 20000 --afterRegionStartLength 3000 (H3K27ac, active enhancers). Signal is then visualised using the plotProfile function.

We determine consensus enhancer and promoter elements by comparing H3K27ac/ H3K4me3-marked regions and consensus mesothelial ATAC-seq peaks. To visualise the accessibility of consensus mesothelial enhancers and promoters, we compute ATAC-seq peak score for each element using the computeMatrix function with following parameters: reference-point --referencePoint center -b 500 -a 500 --missingDataAsZero and visualise using the plotHeatmap function of Deeptools.

Murine H3K27ac, H3K4me3 and H3K27me3 ChIP-seq datasets of E13.5 whole heart were retrieved from the ENCODE project ^109^. Signal p-values were visualised using IGV or Signac where relevant.

We also downloaded the 15-chromatin-state, 8-histone-mark ChromHMM assignments for the mouse E12.5 heart from van de Velde et al. (2021) ^110^.

##### Quantification and statistical analysis

Statistics were performed in R (https://cran.r-project.org/; version 4.4.1) and Python. Differential gene expression analysis was performed using DESeq2, and genes that had BH-adjusted p value < 0.05 was considered to be differential. Statistical significance for differential accessible peaks was calculated using Wilcoxon rank test. To generate chromatin tracks, ATAC-seq data was normalized using the term frequency-inverse document frequency (TF-IDF) method and then visualised either using Signac (version 1.14.0) ^130^ or Integrative Genome Viewer v2.19.5 ^133^. Motif logos were visualised via the ggseqlogo R package (version 0.2) ^172^. Motif locations within open chromatin regions were identified using seqkit ^128^ or Cluster Buster ^129^ and visualised using svist4get ^173^. Heatmap of differential TF binding was made using the EnhancedVolcano R library (version 1.20.0) ^131^.

## Supplementary information titles and legends

**Figure S1. Cadherins profile in the coelomic mesothelium and dynamics of EMT and differentiation in the epicardial lineage**

(A) Genome tracks showing the chromatin accessibility profiles of epicardial cells at E13.5. Track signal is in frequency-inverse document frequency (TF-IDF) normalized counts and merged from N=3 bulk ATAC-seq independent replicates. High-quality peaks were identified by merging replicates and called using MACS2, followed by iterative filtering.

(B) UMAP dimensionality reduction of the mouse lung ATAC-seq dataset (11,638 cells) at E13.5. Mesothelial cells (663 cells) are highlighted by their enriched gene activity of mesothelial markers (*Upk3b* and *Wt1*). Gene activities were imputed based on chromatin accessibility at the relevant gene locus.

(C) UMAP dimensionality reduction of the mouse pancreas ATAC-seq dataset (8,086 cells) at E13.5. Mesothelial cells (750 cells) are highlighted by their enriched gene activity of mesothelial markers (*Upk3b* and *Wt1*).

(D) Matrix plot showing average expression of cadherins across organ mesothelia.

(E) An overview of the analytical workflow used in this manuscript. Created with Biorender.com.

(F) UMAP dimensionality reduction of the integrated epicardial lineage dataset (7,332 cells), combining four datasets profiling four developmental stages from E12.5 to E17.5. Cell type annotations are shown. Arrows indicate direction of differentiation.

(G) Dot-plot showing the mean expression of marker genes and their corresponding cell states in the integrated epicardial lineage scRNA-seq dataset. Dot size represents the fraction of cells in the cell type that express each gene.

(H) Matrix plot showing average expression of cadherins across cell types in our integrated epicardial lineage scRNA-seq dataset.

(I) Plot showing the imputed expression of *Cdh2* over the pseudotime associated with epicardial lineage EMT and differentiation trajectories. An integrated scRNA-seq dataset of lineage-traced epicardial cells, EPDCs, and derivatives in differentiated cell fates was constructed from independent datasets. Branched trajectories associated with EPDC differentiation into specific terminal cell fates are then inferred via pseudotime calculation using Palantir ^139^. Three trajectories are shown, associated with differentiated fates: Pericytes, Cardiac Fibroblasts, and VICs. Plot sections are coloured to represent distinct phases along the EMT and differentiation trajectory, progressing from heart mesothelium/epicardium to EPDC, and subsequently to terminal fates. Vertical lines indicate the boundary pseudotime values demarcating these trajectory phases. MAGIC imputed expression values plotted.

(J) Plot showing the imputed expression of *Cdh3* over the pseudotime associated with epicardial lineage trajectories

(K) Plot showing the imputed expression of *Cdh11* over the pseudotime associated with epicardial lineage trajectories

*EPDC: Epicardium-derived cells; VIC: Valvular interstitial cells; CFb/Fb: Cardiac fibroblast/Fibroblast; SMC: Smooth muscle cell*

**Figure S2. Epigenomic profiling of the developing epicardium**

(A) Profile plots for H3K4me3 histone enrichment scores over gene body regions (±1kb from the transcription start and end sites). Scores were computed using counts per million (CPM)-normalized score of H3K4me3 CUT&RUN-seq generated from the MEC1 epicardial cells.

(B) Profile plots for H3K27ac histone enrichment scores over gene body regions (±1kb from the transcription start and end sites).

(C) Heatmap of normalized peak scores of organ mesothelia in active epicardial enhancer regions, identified using SEACR-called consensus peaks called from H3K27ac CUT&RUN-seq of MEC1 epicardial cells.

(D) Heatmap of normalized peak scores of organ mesothelia in active epicardial promoter regions, identified using SEACR-called consensus peaks called from H3K4me3 CUT&RUN-seq of MEC1 epicardial cells.

(E) Dot-plot showing the percentage of cells and the average scaled expression of co-expression modules across organ mesothelia.

(F) Gene Ontology (GO) term over-representation analysis of genes comprising organ-specific mesothelial modules (heart mesothelium: Meso-M2; lung mesothelium: Meso-M6; pancreas mesothelium: Meso-M3). GO terms were acquired from the GO “Biological Process” database. P- and q-value cutoffs were set at 0.01 and 0.05, respectively.

(G) Module preservation statistics of organ-specific modules Meso-M2, Meso-M3, and Meso-M6 in the corresponding scATAC-seq datasets of the organ mesothelia. Projection of modules to the scATAC-seq datasets were performed by scoring these modules using gene activity computed by chromatin accessibility counts over gene body and promoter regions. The quality of modules and their degree of preservation across modalities were determined through computing Z-summary statistics.

(H) UMAP plot showing the gene activity of *Pitx2* in the E13.5 mouse pancreas scATAC-seq dataset.

(I) UMAP plot showing the gene activity of *Barx1* in the E13.5 mouse pancreas scATAC-seq dataset.

**Figure S3. Epicardial gene regulation by TBX20**

(A) Violin plot showing normalised expression of *Fgf2* in organ mesothelial cells

(B) Genome track showing the proposed gene regulation of the *Sema3d* promoter by a distal putative enhancer (chr3:37398983-37399483; highlighted) linked to TBX20. TF-IDF normalised ATAC-seq tracks show bulk (epicardium) and pseudo-bulk (lung and pancreas mesothelium). Histone tracks show MEC1 epicardial H3K27ac and H3K4me3 CPM-normalized enrichment scores. The link plot shows predicted enhancer-promoter (E-P) connections in the epicardium; link colour denotes the ABC score (the strength of E-P prediction).

(C) Footprint signals across organ mesothelia of the proposed *Fgf2* enhancer. CPM-normalised H3K27ac and H3K4me3 enrichment signal is shown. TOBIAS predicted TBX20 binding site is highlighted.

(D) Violin plot showing the normalised expression of *Sema3d* in organ mesothelial cells

(E) Genome tracks showing the proposed gene regulation of the *Sema3d* promoter by an enhancer (chr5:12507516-12508016; highlighted) linked to TBX20.

(F) Footprint (DNA-binding) signals across organ mesothelia of the proposed *Sema3d* enhancer.

(G) Violin plot showing the normalised expression of *Rarres2* in organ mesothelial cells.

(H) Genome tracks showing the proposed gene regulation of the *Rarres2* promoter by a proximal region linked to TBX20. The CRE is highlighted.

(I) Footprint signals across organ mesothelia of the proposed *Rarres2* CRE.

(J) Violin plot showing the normalised expression of *Pkp2* in organ mesothelial cells.

(K) Genome tracks showing the proposed gene regulation of the *Sema3d* promoter by two enhancers (chr16:16219711-16220211 and chr16:16256606-16257106; highlighted) linked to TBX20.

(L) Normalised ATAC-seq signals of the CRE (chr16:16219711-16220211) from (K) in FACS sorted epicardial cells and EPDCs. Positions of TBX20 binding sites in the enhancer are shown.

(M) Normalised ATAC-seq signals of the CRE (chr16:16256606-16257106) from (K) in FACS sorted epicardial cells and EPDCs. Positions of the TBX20 binding site in the enhancer are shown.

(N) Plot showing the imputed expression of *Pkp2* over the pseudotime associated with epicardial lineage trajectories.

(O) Genome tracks showing the proposed gene regulation of the *Cdh2* promoter by an enhancer (chr18:16740742-16741242; highlighted) linked to TBX20.

(P) Normalised ATAC-seq signals of the CRE (chr18:16740742-16741242) from (O) in FACS sorted epicardial cells and EPDCs. Positions of TBX20 binding sites in the enhancer are shown.

*Epi: Epicardium; EPDC: Epicardium-derived cells; VIC: Valvular interstitial cells; CFb/Fb: Cardiac fibroblast/Fibroblast; SMC: Smooth muscle cells*

**Figure S4. Gene markers for the lung mesothelium are putatively regulated by lung-specific regulons**

(A) Violin plot showing the normalised expression of *Gli1* in E13.5 embryonic lung

(B) Footprint signals across organ mesothelia of the *Hhip* promoter. The exact position of the GLI1 motif is shown. Spearman correlations (rho) between *Gli1*, the predicted bound region, and *Hhip* expression are shown.

(C) Violin plot showing the normalised expression of *Foxf1* in organ mesothelial cells

(D) Genome track showing the proposed gene regulation of the *Pnoc* promoter by a CRE (chr14:65388142-65388642; highlighted) bound by FOXF1. The E-P link predicted by SCENIC+ was illustrated. The FOXF1 binding positions within the CRE are shown.

(E) Violin plot showing the normalised expression of *Foxp2* in organ mesothelial cells

(F) Footprint signals across organ mesothelia of the *Foxp2* promoter. The FOXF1 binding sites in the CRE are shown.

(G) Genome track showing the proposed gene regulation of the *Gli1* promoter by a distal CRE (chr10:127426586-127427086) linked to FOXF1. The CRE is highlighted. The FOXF1 binding positions within the CRE are shown.

(H) Genome track showing the proposed gene regulation of the *Tbx2* promoter by a proximal region linked to FOXF1. The FOXF1-bound region is highlighted. A violin plot showing the normalised expression of *Tbx2* in organ mesothelial cells is shown.

**Figure S5. Gene markers for the pancreas mesothelium are putatively regulated by pancreas-specific regulons**

(A) Genome tracks showing the proposed gene regulation of the *Hoxb8* promoter by a proximal region (highlighted) linked to *BARX1*. The *BARX1* binding site is shown.

(B) Genome track showing the proposed gene regulation of the *Igf1* promoter by a distal region (chr10:87850833-87851333; highlighted) linked to *PITX2*.

(C) Footprint signals across organ mesothelia of the *Cdh6* promoter. The location of PITX2 binding site is shown. The curated PITX2 motif is illustrated.

(D) Violin plot showing the normalised expression of *Isl1* and *Gcg* in organ mesothelial cells

(E) Genome track showing the proposed gene regulation of the *Gcg* promoter by a distal CRE (highlighted) linked to ISL1. The curated ISL1 motif is illustrated and ISL1 binding sites within the CRE are shown.

**Figure S6. Mesothelial CREs drive temporal gene expression**

(A) Stacked violin plot showing the mesothelial median expression of TFs that have motif binding sites in the *Msln* promoter. TFs were identified using the “submerge” function of TOBIAS, which merges all TF binding sites and collects associated footprint information in a region of interest.

(B) Motif logos for WT1 and NRF2/NFE2L2 used in this study.

(C) Plot showing the imputed expression of *Msln* over the pseudotime associated with epicardial lineage trajectories.

(D) Genome tracks showing the regulation of the *Msln* promoter in epicardial EMT and differentiation. The promoter region and two CREs are highlighted. Normalised ATAC-seq tracks of E13.5 epicardial cells and EPDCs are shown. Normalised histone tracks of MEC1 epicardial H3K27ac and H3K4me3.

(E) Violin plot showing the normalised expression of *Tbx20, Fgf2,* and *Csf1* in epicardial cells at E11.5, E13.5, and E17.5.

(F) Genome track showing the regulation of the *Csf1* promoter in epicardial cells over embryonic development. Merged normalised ATAC-seq tracks of E11.5, E13.5, and E17.5epicardial cells (N = 3 per stage). Normalised histone tracks show MEC1 epicardial H3K27ac and H3K4me3. The enhancer predicted to regulate *Csf1* is highlighted (Refer to Figure 3H).

(G) Genome tracks showing the regulation of the *Fgf2* promoter in epicardial cells over embryonic development. The enhancer predicted to regulate *Fgf2* is highlighted (Refer to Figure S3B).

(H) Violin plot showing the normalised expression of *Barx1 and Hoxb8* in pancreas mesothelial cells at E12.5, E13.5, E14.5, and E17.5.

(I) Genome tracks showing the regulation of the *Hoxb8* promoter in the pancreas mesothelium over embryonic development. Normalised ATAC-seq signal tracks of E13.5, E14.5, and E17.5 pancreas mesothelial cells are shown. The enhancer predicted to regulate *Hoxb8* is highlighted (Refer to Figure S5A).

(J) Violin plot showing the normalised expression of *Isl1 and Gcg* in pancreas mesothelial cells at E12.5, E13.5, E14.5, and E17.5.

(K) Genome tracks showing the regulation of the *Gcg* promoter in the pancreas mesothelium over embryonic development. The enhancer predicted to regulate *Gcg* is highlighted (Refer to Figure S5E).

*Epi: Epicardium*

**Figure S7. Inference of cardiac cell-type specific gene regulatory networks**

(A) UMAP dimensionality reduction of the single-cell E13.5 mouse heart ATAC-seq dataset (7,292 cells). CM (2,262 cells) are highlighted by the black arrow, inferred by *Myl2* gene activity. Gene activities were imputed based on chromatin accessibility at the relevant gene locus.

(B) UMAP dimensionality reduction of the single-cell E13.5 mouse heart ATAC-seq dataset. Cardiac vascular ECs (216 cells) are highlighted by the black arrow, inferred by *Fabp4* gene activity.

(C) Correlation matrix of E13.5 cardiac cell types and organ mesothelia based on chromatin-accessible peak signatures.

(D) Upset plot showing the intersection of chromatin peaks from the E13.5 mesothelium across the heart, lung, and pancreas and E13.5 cardiac cell types (CM and CEC).

(E) Pearson correlation matrix of E13.5 cardiac cell types and three organ mesothelia based on transcriptomic signatures. Colour is based on correlation coefficient values (higher value indicates stronger positive correlation)

(F) Dot-plot showing the expression of marker genes and their corresponding cell types. Cells are grouped based on their similarities in a principal component analysis (PCA) representation using a dendrogram.

(G) Motif logos of MAF and TCF21.

(H) Pairwise comparison of TF activity between the heart mesothelium and vascular ECs. The volcano plots show the differential binding activity against the −log10(p value) of all investigated TF motifs; each dot represents one motif. Only motifs for TFs expressed in at least one organ mesothelium are retained. Organ-mesothelium-specific TFs are labelled in red.

(I) Pairwise comparison of TF activity between the heart mesothelium and ventricular CM.

*(C)EC: (cardiac) endothelial cells; CM: cardiomyocytes*

**Figure S8. Conserved CREs regulate mesothelial gene marker expression**

(A) Violin plot showing the normalised expression of *Tcf21* and *Maf* in mesothelial cells and cardiac cell types.

(B) Stacked violin plot showing the median expression of *Maf, Krt8, Krt18,* and *Upk3b* in epicardial cells at E11.5, E13.5, and E17.5. Higher expression is shown in yellow.

(C) Genome tracks showing the regulation of the *Krt18* promoter in epicardial cells over embryonic development. The enhancer predicted to regulate *Krt18* is highlighted (Refer to Figure 6B).

(D) Genome tracks showing the regulation of the *Krt8* promoter in epicardial cells over embryonic development. The enhancer predicted to regulate *Krt8* is highlighted (Refer to Figure 6E).

(E) Genome tracks showing the regulation of the *Upk3b* promoter in the epicardium over embryonic development. The enhancer predicted to regulate *Upk3b* is highlighted (refer to Figure 6M).

(F) Stacked violin plot showing the median expression of *Maf, Krt8, Krt18,* and *Upk3b* in pancreas mesothelial cells at E12.5, E13.5, E14.5, and E17.5. Higher expression is shown in yellow.

(G) Genome tracks showing the regulation of the *Krt18* promoter in the pancreas mesothelium over embryonic development. The *Krt18*-linked enhancer and *Krt18* promoter are highlighted.

(H) Genome tracks showing the regulation of the *Krt8* promoter in the pancreas mesothelium over embryonic development. The *Krt8*-linked enhancer and *Krt8* promoter are highlighted.

(I) Genome tracks showing the regulation of the *Upk3b* promoter in the pancreas mesothelium over embryonic development. The *Upk3b*-linked enhancer and the *Upk3b* promoter are highlighted.

(A) (L) Stacked violin plot showing the median expression of *Maf, Krt8, Krt18,* and *Upk3b* in lung mesothelial cells at E12.5, E13.5, E15.5, E17.5, and P3. Higher expression is shown in yellow.

(B) (G) Genome track showing the regulation of the *Krt18* promoter in the lung mesothelium over embryonic development.

(C) (H) Genome track showing the regulation of the *Krt8* promoter in the lung mesothelium over embryonic development.

(J) Genome track showing the regulation of the *Upk3b* promoter in the lung mesothelium over embryonic development.

**Figure S9. Mouse-Human comparison of the embryonic epicardium**

(A) UMAP dimensionality reduction of the cross-species integrated epicardial cells labelled by species.

(B) UMAP plot of the cross-species integrated epicardial cells labelled by the developmental stage

(C) Stacked violin plot showing the median expression of marker genes that demarcate epicardial cell states in the cross-species dataset. Higher expression is shown in yellow.

(D) Heatmap showing the enrichment of GO terms in epicardial cell states. Depth of colours indicates AUCell enrichment score (darker colour indicates higher score).

(E) Violin plot showing the normalised expression of *Cdh1* between species in the cross-species integrated epicardial dataset.

(F) UMAP plot showing the cell types in mouse embryonic and postnatal heart integrated dataset (72,120 cells), spanning early development (E10.5 to P6).

(G) Violin plot showing the expression of *Cdh1* in mouse embryonic and postnatal heart integrated dataset.

(H) Violin plot showing the expression of *Cdh1* in a published integrated mouse embryonic heart dataset ^82^.

(I) Genome tracks showing the *Cdh1* promoter and proposing evidence for PRC2-mediated H3K27me3 deposition as a mechanism that silences the expression of this gene in the mouse heart. Track signals include normalised ATAC-seq profiles of organ mesothelia and cardiac cell types; Normalized MEC1 epicardial H3K4me3 enrichment score; Normalized H3K4me3 enrichment score of the mouse whole heart at E13.5 from the ENCODE project; Normalized H3K27me3 enrichment score of the mouse whole heart at E13.5 from the ENCODE project; Mouse embryonic heart ChromHMM-based model with 9 histone marks and 15 states. The ChromHMM model was used to annotate the *Cdh1* promoter.

*EC: Endothelial cell; CM: Cardiomyocyte*

**Supplementary Table 12.**
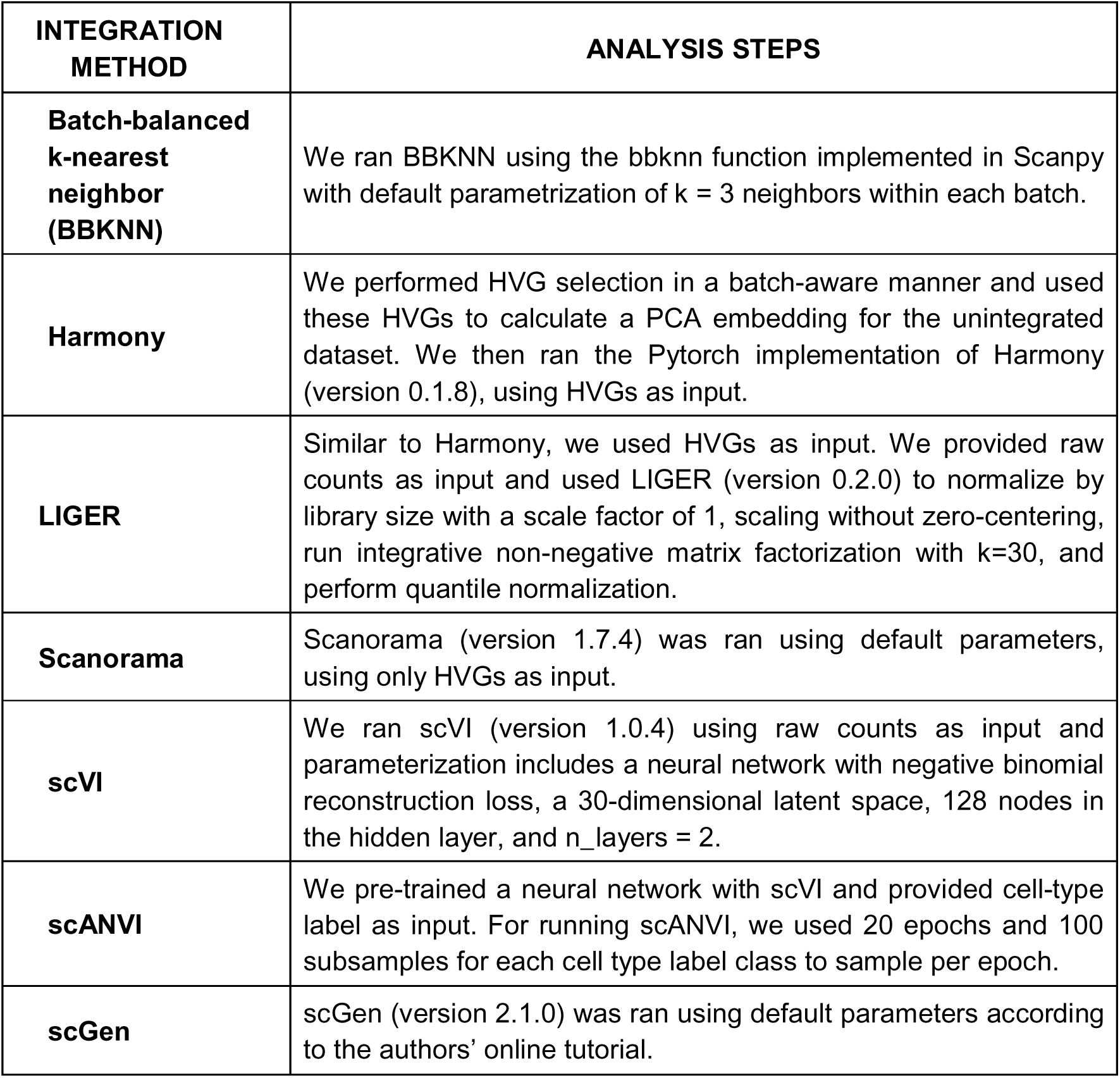
Analysis steps and parameterization used for single-cell integration algorithms used in this study.

| **INTEGRATION METHOD** | **ANALYSIS STEPS** |
| --- | --- |
| **Batch-balanced k-nearest neighbor (BBKNN)** | We ran BBKNN using the bbknn function implemented in Scanpy with default parametrization of $k = 3$ neighbors within each batch. |
| **Harmony** | We performed HVG selection in a batch-aware manner and used these HVGs to calculate a PCA embedding for the unintegrated dataset. We then ran the Pytorch implementation of Harmony (version 0.1.8), using HVGs as input. |
| **LIGER** | Similar to Harmony, we used HVGs as input. We provided raw counts as input and used LIGER (version 0.2.0) to normalize by library size with a scale factor of 1, scaling without zero-centering, run integrative non-negative matrix factorization with $k=30$, and perform quantile normalization. |
| **Scanorama** | Scanorama (version 1.7.4) was ran using default parameters, using only HVGs as input. |
| **scVI** | We ran scVI (version 1.0.4) using raw counts as input and parameterization includes a neural network with negative binomial reconstruction loss, a 30-dimensional latent space, 128 nodes in the hidden layer, and $n\_layers = 2$. |
| **scANVI** | We pre-trained a neural network with scVI and provided cell-type label as input. For running scANVI, we used 20 epochs and 100 subsamples for each cell type label class to sample per epoch. |
| **scGen** | scGen (version 2.1.0) was ran using default parameters according to the authors' online tutorial. |

## Notes

### Competing Interest Statement

The authors have declared no competing interest.

### Summary of Updates

In this revised version, the figure files accompanying the manuscript, including supplementary figures, have been improved. Specifically, the files were reformatted in accordance with the guidelines to improve file resolution and quality.

